# Domestication, spread, and improvement have shaped biological nitrogen fixation traits in *Zea* taxa

**DOI:** 10.64898/2026.09.30.755791

**Authors:** Keyvan Maleki, Layne Connolly, Nicola Lorenz, Tuny Amphonechit, Alexander J. Lindsey, Hugo Perales, Vivian Bernau, Richard Dick, Kristin Mercer

## Abstract

- Synthetic nitrogen fertilizers enhance maize (*Zea mays* ssp*. mays*) productivity but impose substantial environmental and economic costs. Some Mexican maize landraces support biological nitrogen fixation (BNF) in aerial root mucilage, raising the question of how much variation exists in this trait and how domestication, geographic spread, and improvement have shaped BNF-related traits across *Zea* in the Americas.
- We evaluated aerial root traits and BNF capacity across the maize domestication continuum using 60 *Zea* accessions grown in field experiments over two years under low-N conditions.
- Some accessions of wild *Zea*, landraces from Mexico and South America, and improved varieties had elevated nitrogen derived from the atmosphere (%Ndfa), ranging from 20 to 40%; some of these also had greater %Ndfa than typical improved varieties. Spread and improvement appear to have significantly reduced the number of aerial root-bearing nodes, with wild *Zea* and Mexican landraces consistently producing more. Nevertheless, the proportion of nitrogen derived from the atmosphere (%Ndfa) does not vary systematically across the domestication continuum.
- Thus, crop evolution shaped aerial root phenotypes more than associative BNF capacity. The persistence of substantial within-group variation indicates that the genetic potential for enhanced BNF remains within the *Zea* gene pool, providing opportunities to encourage and improve nitrogen use efficiency in maize cropping systems.

## Introduction

Nitrogen (N) is a critical nutrient limiting crop yield, and modern agriculture relies heavily on synthetic N fertilizers to sustain productivity (Zhang *et al*., 2015; Ladha *et al*., 2016). In cultivated maize (*Zea mays* ssp. *mays*), one of the world’s most important cereals, large fertilizer inputs are routinely applied to maintain yields (Baum *et al*., 2025). Yet, NUE of applied N fertilizer remains low, with substantial portions being lost through leaching, surface runoff, and gaseous emissions (Govindasamy *et al*., 2023; Martinez-Feria *et al*., 2024). These inefficiencies impose major ecological and economic costs: nitrate pollution degrades waterways; nitrous oxide (N_2_O), a potent greenhouse gas, exacerbates climate change; and farmers face escalating fertilizer expenses (Bloch *et al*., 2020; Hassan *et al*., 2022; Bhatt *et al*., 2025). Reducing reliance on synthetic fertilizers and improving NUE are therefore urgent priorities for sustainable agriculture.

Harnessing biological nitrogen fixation (BNF), the microbial conversion of atmospheric N_2_ into plant-available forms, has long been viewed as a major goal for cereal improvement (Triplett, 1996). Whereas legumes meet much of their N demand through symbioses with rhizobia in root nodules, maize and other cereals lack such specialized structures (Rosenblueth *et al*., 2018; Pankievicz *et al*., 2019; Ladha *et al*., 2022). Instead, grasses associate with free-living, endophytic, or rhizosphere diazotrophs that can contribute fixed N in low-input systems (Gupta *et al*., 2019; Guo *et al*., 2023). For several decades now, maize roots and tissues have been shown to harbor diazotrophic bacteria. Early studies using ^15^N isotope dilution and enrichment approaches, together with culture-based surveys, demonstrated that maize can associate with N-fixing taxa such as *Azospirillum, Herbaspirillum*, *Pantoea*, and related genera. These studies provide evidence for BNF activity in maize roots and rhizospheres, although the magnitude of plant N gain has remained uncertain (Chelius & Triplett, 2001; Roesch *et al*., 2008; Montañez *et al*., 2009). Most studies confirm low rates of BNF in the maize rhizosphere under low N conditions (Goldstein & White, 2022; Connolly *et al*., 2025; Zhao *et al*., 2025). Furthermore, high N fertilizer rates suppress nitrogenase activity (Wen et al., 2021), making it challenging to optimize this BNF to meet the high N demand of maize (Guo et al., 2023). Biological N fixation rarely supplies a significant amount of N to meet the needs of cereals, with the exception of sugarcane (*Saccharum officinarum* (Urquiaga *et al*., 2012), rice (Boddey *et al*., 1995), and wheat (Iniguez *et al*., 2004). Synthetic-biology approaches may show promise: *Klebsiella varicola* strains with nitrogenase uncoupled from exogenous N sustained fixation under fertilized conditions and produced modest yield gains (Wen *et al*., 2021; Pankievicz *et al*., 2022).

Recent discoveries in maize have shown that certain landraces can sustain substantial BNF through specialized aerial roots and their mucilage secretions (Van Deynze *et al*., 2018; Bennett *et al*., 2020; Pankievicz *et al*., 2022; Gao *et al*., 2023; Connolly *et al*., 2025; Zhao *et al*., 2025; Laspisa *et al*., 2025). The Sierra Mixe landrace of southern Mexico, for example, produces many tiers of aerial roots that secrete sugar-rich mucilage under high-moisture conditions (van Deynze *et al*., 2018). This mucilage forms a carbon-rich, low-oxygen microenvironment that harbors diazotrophic taxa and *nif* genes, supporting active nitrogenase activity verified by acetylene reduction and ¹ N incorporation assays (Bennet *et al*., 2020; Gao *et al*., 2023). Field trials indicate that BNF can supply 29–82% of plant N in this landrace (van Deynze *et al*., 2018). Importantly, this capacity is not unique to Sierra Mixe maize as other Mexican landraces also support diazotrophs and benefit from BNF (Gao *et al*., 2023; Connolly *et al*., 2025). Moreover, variation in aerial root traits (number, diameter, and mucilage viscosity) can strongly correlate with nitrogenase activity (Pankievicz *et al*., 2022; Gao *et al*., 2023; Venado *et al*., 2025).

Yet the evolutionary origin of these associations remains unresolved. Maize (*Zea mays* ssp. *mays*) was domesticated from the lowland teosinte *Z. mays* ssp. *parviglumis* in southwestern Mexico and subsequently experienced introgression from the highland teosinte *Z. mays* ssp. *mexicana* as it spread into the Mexican highlands (Yang et al., 2017).There was subsequent spread of maize throughout what is now Mexico, Latin America and the US, resulting in diversification from both hybridization and natural/farmer-mediated selection (Muntean *et al*., 2022; Yang *et al*., 2023; Gutaker, & Purugganan, 2024). Modern breeding has further contributed to evolutionary change (Cahn *et al*., 2024). Evidence from *Z. mays* ssp*. mexicana* shows that its mucilage harbors diazotrophic activity, suggesting an ancient association that may have been retained or amplified in some maize lineages (Gao *et al*., 2023). The domestication gradient from teosinte through traditional landraces to improved varieties, offers a unique framework to disentangle how evolutionary forces, farmer selection, and modern breeding have shaped traits like aerial root development, mucilage secretion, and diazotrophic associations (Hufford *et al*., 2012; Swarts *et al*., 2017; Janzen *et al*., 2022). While shifts in root architecture, microbial interactions, and nutrient acquisition strategies across this gradient are documented (Kremling *et al*., 2018; Wallace *et al*., 2018; Emmet *at al.,* 2022), the extent to which associative N-fixation traits were retained, reduced, or lost remains unclear. Emerging evidence suggests that domestication and breeding may have eroded cereal’s capacity for associative N fixation (Yue *et al*., 2023; Guo *et al*., 2023; Chang *et al*., 2025). This pattern is consistent with long-standing agroecological evidence showing that crop intensification and reliance on synthetic N has reduced biologically mediated N inputs and microbial contributions to plant N nutrition (Drinkwater *et al*., 1998; Drinkwater & Snapp, 2007; Isaac *et al*., 2021; He *et al*., 2025). Yet, systematic comparisons across a domestication gradient, including teosinte, landraces, and improved varieties, are lacking.

The ^15^N natural abundance approach provides a means to evaluate atmospheric N fixation under field conditions without artificially enriching the soil or plant environment with isotopic tracers. The approach exploits differences in isotopic composition between atmospheric N_2_ and soil-derived N: atmospheric N_2_ has a relatively constant δ^15^N value near 0‰, whereas soil N pools often become enriched in ^15^N through fractionating processes such as mineralization, nitrification, and denitrification (Shearer & Kohl, 1986; Högberg, 1997; Boddey et al., 2000; Unkovich et al., 2008; Cox et al., 2022; Pang et al., 2023). Consequently, when soil N sources are sufficiently enriched relative to atmospheric N_2_, plants obtaining a greater proportion of their N through biological N fixation are expected to exhibit lower δ^15^N values than appropriate non-N_2_-fixing reference plants. Comparisons of plant and reference δ^15^N values can therefore be used to estimate the proportion of plant N derived from the atmosphere (%Ndfa), providing a field-based approach for comparing atmospheric N fixation among genotypes or germplasm groups.

Here, we address this gap by evaluating a broad representation of maize germplasm from across the Americas for traits related to biological N fixation in *Zea* taxa. Particularly, this study aimed to understand how crop evolution, including domestication, geographic spread, and modern improvement, has influenced aerial root development and N fixation traits in *Zea* under field conditions. More specifically, we ask: (i) how much variation exists among Zea accessions in aerial root development, mucilage production, plant growth, and N derived from the atmosphere; and (ii) how have domestication, geographic spread, and modern improvement shaped these BNF-related phenotypic traits and N derived from the atmosphere?

## Materials and Methods

### Plant material and accession selection

To assess the effect of the domestication, spread, and improvement (see Fig. 1) on BNF related traits and BNF rates in *Zea* taxa, we selected 58 maize accessions, including wild teosinte (n = 8), Mexican landraces (n = 20), South American landraces (n = 11), U.S. landraces (n = 4), heirloom varieties (n = 5), and improved hybrids/varieties (n = 10). For isotopic comparisons, we used four improved maize varieties (Painted Mountain and LAT, Lamb 333, FOS 8500) from this panel as an internal non-N-fixing reference and included two additional non-maize reference species (sunflower and sorghum Sudangrass). (Table S1). To measure N derived from the atmosphere (%Ndfa), these three reference plants were used as a standard against which to compare isotopic levels of *Zea* accessions (see BNF measurement section for further detail).

**Figure 1.**
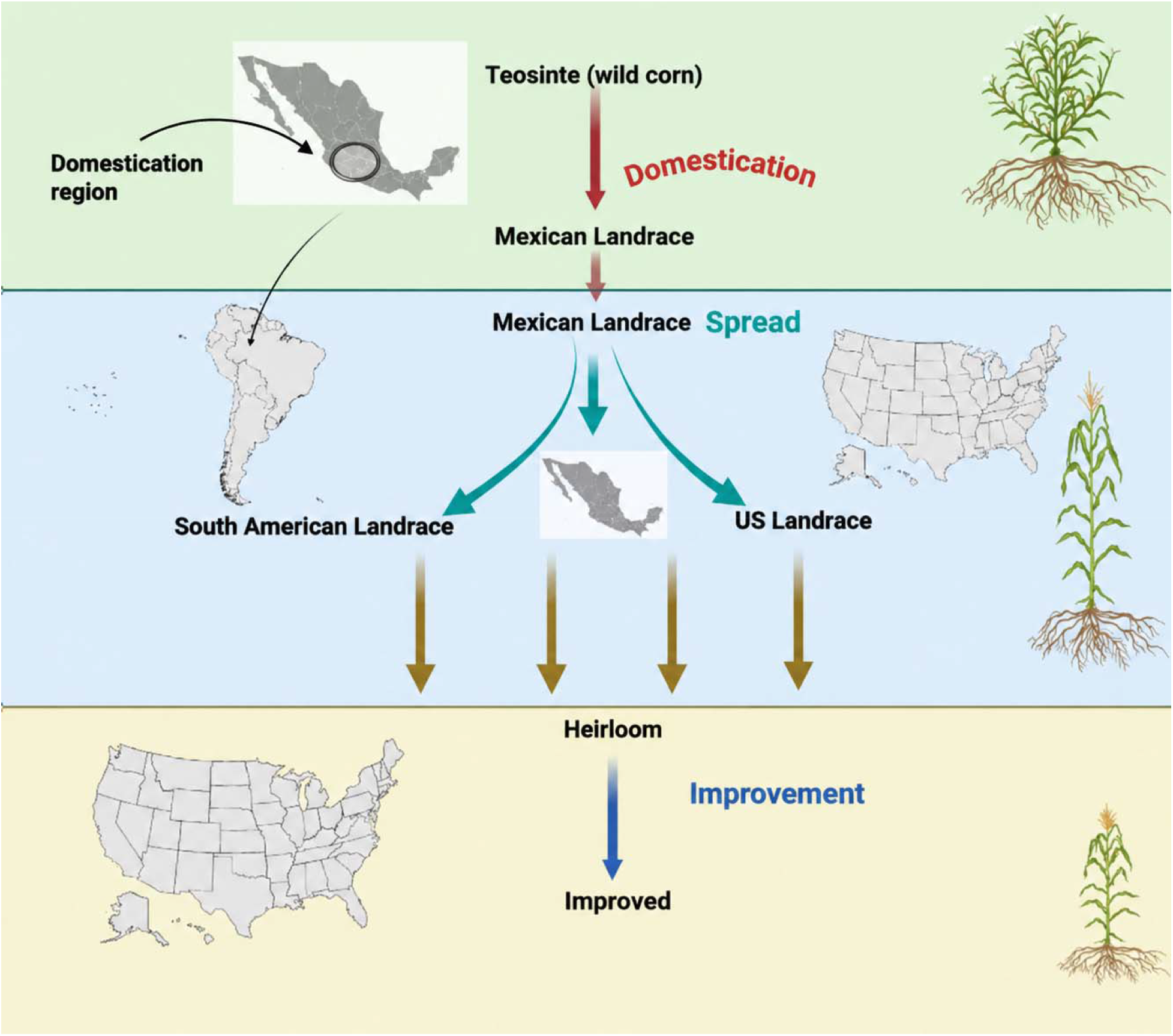
Schematic illustration of the domestication, geographic spread, and improvement of maize (*Zea mays* ssp. *mays)*. Teosinte, the wild ancestor of maize, was domesticated in Mexico, giving rise to landraces. From there, maize spread within Mexico, as well as southward to South America and northward to the United States, where distinct landrace groups developed. Subsequent breeding and selection led to heirloom varieties and, more recently, modern improved cultivars. The arrows indicate the transitions of domestication, spread, and improvement that shaped maize diversity.

To curate this comprehensive list of accessions across the domestication gradient (Table S1), along with associated location and race information, we utilized the USDA-ARS National Plant Germplasm System (NPGS) Germplasm Resources Information Network (GRIN; https://npgsweb.ars-grin.gov/gringlobal/search). The National Plant Germplasm Systems (NPGS) GRIN-Global database provided extensive records on accessions, including details on race, name, date, elevation, and geographic coordinates of the collection location (https://npgsweb.ars-grin.gov/gringlobal/search). We then refined our record list to accessions with available seeds and chose accessions that represented the geographic range (e.g., Mexico, South America, and US for landraces), the range of racial categories for landraces, the range of species and subspecies for wild *Zea* taxa, and other known phenotypic variation. For teosintes and Mexican and South American landraces, we consulted recent studies describing diversity in maize root architecture and aerial root morphology to identify accessions with contrasting root phenotypes (Hostetler et al., 2022; 2026). We also reviewed studies on biological nitrogen fixation (BNF) in heirloom and improved maize varieties to include accessions for which BNF had previously been reported, such as Hickory King (Van Deynze et al., 2018). We categorized accessions (wild and landrace) into three elevational groups: lowland (<1000 m), midland (1000–2000 m), and highland (>2000 m); our main objective for elevation was to have a good representation of elevations as maize has shown local adaptation in response to elevation, and environmental conditions (Mercer *et al*., 2008; Janzen *et al*., 2022). For heirloom and improved maize, accessions were selected to represent diverse agronomic backgrounds, including heirloom varieties available in Ohio, improved hybrids widely used as research standards (e.g., B73 × Mo17), and varieties previously identified as contrasting in nitrogen-use efficiency (NUE; see Table S1; Goldstein., et al., 2019; Goldstein, 2022; Goldstein & White, 2026) (C2.B2.C46, FOS 8500, Lamb333, and LAT7-113 x LH123-13).

### Field experiment

To study the effect of domestication, spread, and improvement on the ability of maize to fix atmospheric N, two field experiments (one in the summer of 2023 and the other in the summer of 2024) were performed at the Ohio State University Waterman Agricultural and Natural Resources Laboratory in Columbus, Ohio (40°00’32.4"N 83°02’14.5"W; 226 masl; silt loam soil). In both years, our experimental design was a randomized complete block framework, incorporating four blocks to ensure statistical robustness and to account for environmental variability. A very small amount of N fertilizer (ammonium polyphosphate - 10 kg N ha^-1^) was applied at the V4-V6 stage of maize in two doses. Each maize accession was planted in randomized plots with rows spaced 0.75 m apart and plants spaced 0.33 m apart within each row. Each row within each plot contained seven plants, except for teosinte, for which five plants were included per plot to accommodate its larger growth form.

In this experiment, an irrigation regime was implemented to induce mucilage production on aerial roots at nodes along the stem. The irrigation system, equipped with sprinklers and risers (generic riser tubing + PVC), was designed to maintain a high relative humidity within the canopy to help facilitate mucilage production, rather than primarily for the purpose of watering the plants. The irrigation schedule consisted of five daily sessions, each lasting five minutes. The timing changed to 10 minutes in summer 2024 during a droughty period. This frequency and duration were chosen to increase relative humidity without over-saturating the plants. Based on standard rainfall reporting metrics, the output of our irrigation system was set to deliver 2.2 L/m² in each 5-minute session. This equates to a rate of approximately 26.4 mm/hr or 12 mm per day over the course of the five applications.

### Data collection: plant phenotyping

We measured a range of traits associated with plant performance and related to N fixation on aerial roots. Mucilage rating was evaluated through a qualitative assessment using a predefined scale designed to capture both the amount and consistency of mucilage produced by each plant. Mucilage production was rated on a scale from 0 to 3, where a rating of 0 indicates absence of mucilage and 3 signifies abundant mucilage visibly dripping from aerial roots (Fig. S1). Observations were conducted once per week throughout the growing season and additionally following rainfall events, as increased relative humidity promotes mucilage secretion. Assessments were focused on younger aerial roots, which are more likely to exude mucilage than older nodes (Pankievicz *et al*., 2022). For data analysis, ordinal observation of mucilage presence was used (0-3). Other traits assessed included number of layers of nodes with aerial root (ARLN), diameter of aerial roots (mm), length of the first and second internodes (cm), stem diameter (mm) and plant height (cm). Plants were harvested at VT-R2 stages. To take data on each trait, we cut down three plants per plot marked for phenological stage. A more comprehensive description of data collection can be found in the supplementary file (trait measurement section).

### Nitrogen isotopic measurements

To assess actual N fixation under field conditions, the ^15^N natural abundance method was applied to all 60 accessions (Van Deynze *et al*., 2018). Whole-plant samples (shoot tissues; above ground materials including aerial roots) from up to three plants per plot were processed using a woodchipper (model: SC-3013, 3-inch chipper shredder; ECHO Incorporated). We then subsampled the biomass, oven-dried at 55°C to constant mass, and ground to a fine 100-mesh powder using a Udy cyclone mill (UDY Corporation, CO). The mill and woodchipper were cleaned thoroughly between samples to prevent cross-contamination. The mill was cleaned using a vacuum, compressed air, and 80% ethanol, whereas the woodchipper was cleaned with pressurized water only. For N analysis, approximately 5 mg of dried plant tissue was weighed into tin capsules and analyzed using an elemental analyzer (CHN EA 1108, Carlo Erba Instruments, Italy) coupled to a Delta V Advantage Isotope Ratio Mass Spectrometer (IRMS). The δ^15^N values were expressed in per mil (%) relative to atmospheric N, calculated as (Coplen *et al*., 2011):

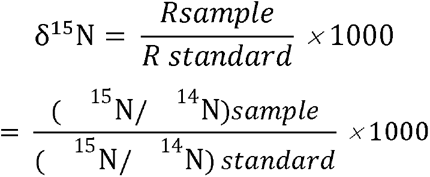

where R represents the ^15^N/^14^N ratio. Atmospheric N [δ^15^N (R_standard_)= 0.00366295] served as the standard, and acetanilide was used for calibration. The proportion of N derived from atmospheric fixation (%Ndfa) was estimated as (Boddey *et al*., 2000; Sarr *et al*., 2016):

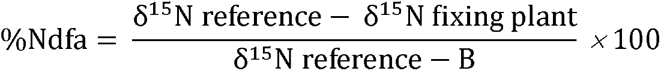

where B represents the δ^15^N abundance in atmospheric N (assumed to be zero) (Unkovich *et al*., 2008). We used sunflower, sorghum Sudangrass, and four improved maize varieties (Painted Mountain and LAT, Lamb 333, FOS 8500) as putatively non-fixing reference plants. Sunflower and sorghum Sudangrass were selected because they are non–N-fixing species (Van Deynze *et al*., 2018; Venado *et al*., 2025). To estimate %Ndfa, multiple reference plants were used. In addition, we calculated %Ndfa based on the average of all reference plants. The maize varieties used for reference had consistently showed low δ¹ N values across two years in the field and within the literature (Goldstein & White, 2020; Connolly *et al*., 2025). Total aboveground N accumulation (kg N ha ¹) was calculated as the product of aboveground dry biomass (kg ha ¹) and tissue N content (%). The amount of fixed N (kg/ha) was estimated by multiplying %Ndfa by the N concentration and total aboveground biomass. (Barros *et al*., 2020)

### Statistical approach

For all analyses of phenotypic traits and N isotope variables, the generalized linear mixed models (GLMMs) analysis was done with the *lme4,* and *lmerTest,* and *emmeans* packages in R (version 4.4.1; R Core Team, 2024). To confirm the results obtained from R, analysis was also done using SAS (SAS Institute Inc. version 9.4) We ran two models on each of our response variables: one to elucidate difference among domestication levels and one to explore differences among individual accessions. For the first, domestication was treated as a fixed effect [domestication levels included wild non-progenitors, wild progenitors, Mexican landraces, South American landraces, US landraces, heirloom, and improved varieties], while accession, block, and the accession-by-block interaction were included as random effects to account for subsampling and experimental structure. Each year was analyzed separately. For the second model, accession was treated as a fixed effect, while block and the block-by-accession interaction were included as random effects to account for spatial variation and subsampling within accession-by-block plots, respectively. Each year was analyzed separately. Alternative response variable distributions were evaluated using Akaike’s Information Criterion (AIC) and log-likelihood tests. For number of nodes bearing aerial roots, aerial root diameter, and mucilage, we modeled the response variable using a negative binomial distribution with a log-link function rather than a normal distribution. This provided the best fit and appropriately accounted for overdispersion in these discrete counts.

Degrees of freedom were approximated using the Kenward-Roger method. To assess differences between domestication categories and accessions, least squares means (LSMeans) were estimated and compared with Tukey adjustments for multiple testing. To assess the significance of random effects, a Wald chi-square and log-likelihood tests were performed on the covariance parameters.

We performed two additional tests. First, we tested whether the estimated marginal mean (least-squares mean) of %Ndfa for each accession differed significantly from zero using contrasts of model-estimated marginal means implemented with the *emmeans* package in R (Lenth, 2023).Second, to evaluate whether maize varieties commonly cultivated in the U.S. Midwest (i.e., O-6160, 107-day and SX-649) differed in biological nitrogen fixation (BNF) capacity compared with the other Zea accessions, we performed a Dunnett’s test to compare multiple treatment groups against a predefined control while controlling the family-wise Type I error rate (Bretz et al., 2016). Here, the typical Midwest-grown improved maize varieties served as the control group because they are most like those used in conventional U.S. production systems. Least-squares means estimated from the fitted generalized linear mixed models were compared between each accession and the Midwest control group.

## Results

### Variation in nitrogen-fixing traits among all accessions

Accessions varied for traits related to aerial root development, growth, and BNF across two years (Figs 2, 3 and S2-6; Tables 1, S2, and S3). The number of nodes with aerial roots (ARLN) varied from around 12 to less than 1 in 2023 and from more than 15 to around 0 in 2024, though many accessions were not statistically distinguishable from one another due to high variation in expression within accessions (Fig. 2). Nevertheless, wild teosintes and Mexican and South American landrace accessions topped the list for highest estimates of ARLN (Fig. 2). Notably, the highest ARLN estimates in both years were observed a wild crop-progenitor (*Z. mays* ssp. *mexicana*; PI 658195) and a Mixeño landrace from Oaxaca, Mexico (Ames 19897); improved and heirloom accessions tended to have lower values for ARLN (Fig 2).

**Figure 2.**
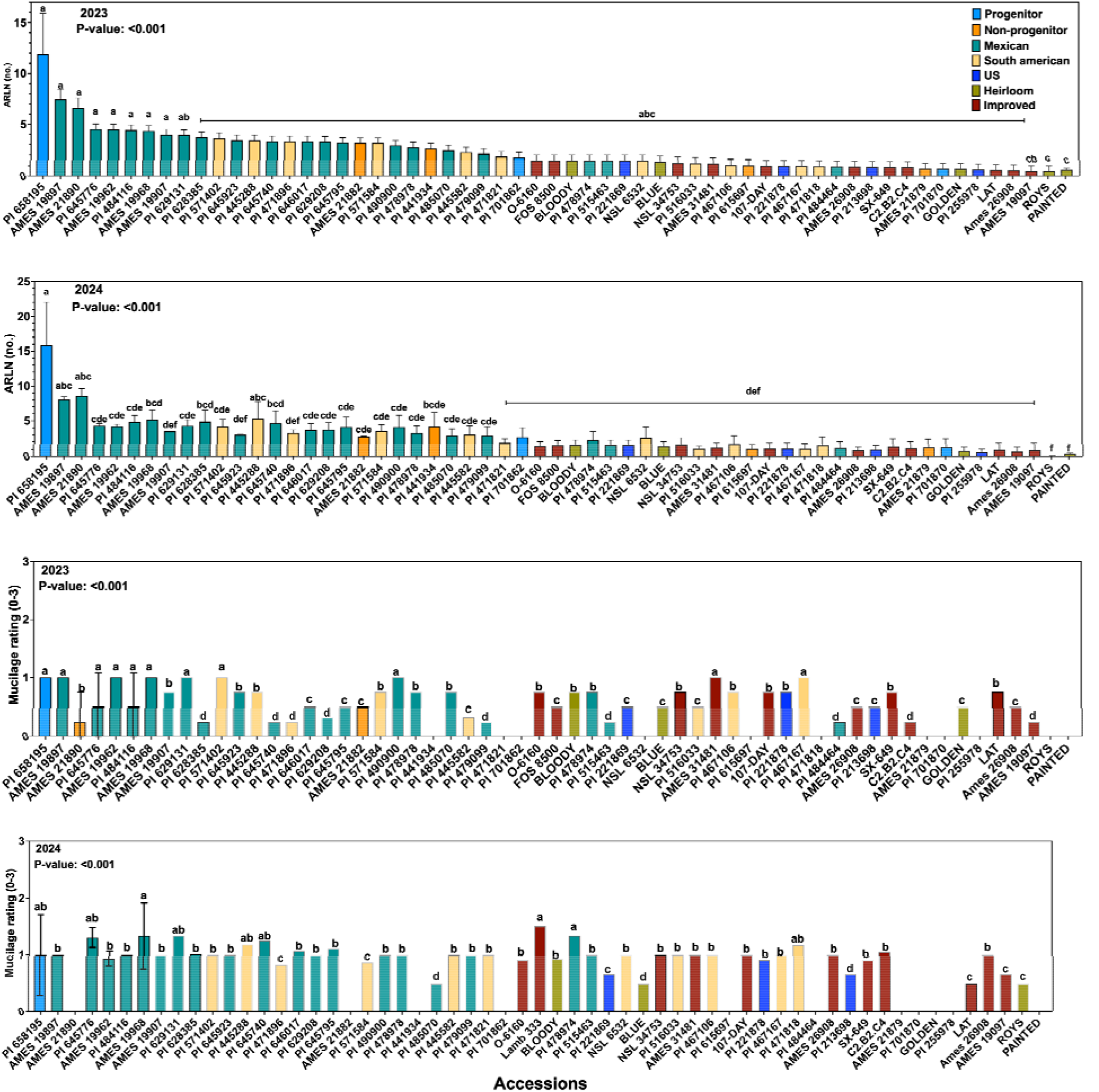
Variation in number of nodes with aerial roots (ARLN) and mucilage across diverse *Zea* accessions and domestication gradient in 2023 and 2024. P-values denote the effect of accession on ARLN and mucilage. P-values were derived using generalized linear mixed models (GLMMs). The absence of error bars reflects either missing data/replicates or identical mucilage scores among replicates. Means that share a letter are not significantly different (Tukey’s adjustment, P < 0.001).

Mucilage production also varied significantly among accessions (Fig 2; Table S2). In 2023, most accessions exhibited relatively low mucilage rating averages (ranging from 0 to 1 on a three-point scale in 2023 and 0 to ∼1.5 in 2024). The low scores in both years likely reflect the intermediate humidity levels in Ohio; the substantially drier conditions experienced in 2023 may have reduced mucilage further. Several accessions exhibited increased mucilage production in 2024 compared to 2023, while others remained consistently low, e.g., two Mexican landraces, AMES 19968 (Bolita) and PI 645776 (Ancho; Fig 2).

Traits indicative of size varied among accessions (Tables S2, S3; Fig. S2-6). Notably, some wild teosinte and Mexican landrace accessions [e.g. PI 658195 (crop progenitor, *Z. mays* ssp. *mexicana*), PI 645923 (Jala landrace from Nayarit, Mexico)] produced the tallest plants in both years (Fig. S2-6). Other traits, including internode lengths and aerial root diameters, showed more variable and inconsistent patterns across years suggesting greater sensitivity to environmental conditions or developmental timing. Mexican landraces consistently exhibited the thickest stem diameters among all domestication groups, particularly accessions PI 645740 (a Comiteco landrace from Chiapas, Mexico) and PI 645923 (a Jala landrace from Nayarit, Mexico) (Fig. S2-6). Wild teosinte (both progenitor and non-progenitor) accessions had the highest plant biomass (50000 kg/h^-1^ approximately) and heirloom and improved varieties had the lowest (1000 kg/h^-1^) (Fig S7).

Total N content (%) varied significantly among accessions in both years, ranging from 2% to 0.2% (Table 1, Fig S8). In 2023, most accessions showed moderate levels of total N; some accessions, such as PI 516033 (0.9 %; a South American landrace from Argentina) and PI 484464 (1%; a Cristalino de Chihuahua from Chihuahua, Mexico), exhibited higher total N content while others, such as PI 645795 (0.4%; a Celaya landrace from Guanajuato, Mexico) and PI 645776 (0.3%; an Ancho landrace from Guerrero, Mexico), had lower N content (Fig S8). In 2024, variation in total N content was more pronounced. Improved varieties LAT (∼1.3%) and Lamb 333 (∼1.5%; both putatively non-fixing improved varieties) and the heirlooms Painted Mountain (a putatively non-fixing variety based on our previous work and confirmed across both growing seasons; ∼1.4%) and Roy’s Calais Flint (∼1.3%) exhibited elevated N content, while others such as PI 645923 (0.4%; the Jala landrace from Nayarit, Mexico) and PI 629131 (0.3%; a Tuxpeño landrace from Chiapas, Mexico) showed lower values. δ¹ N-based estimates of biological N fixation showed no significant accession-level effect in either year (Table 1), although averages varied from 0 to 15 in both years. Similarly, percent N derived from the atmosphere (%Ndfa) did not significantly differ among accessions (Table 1; Fig. 3). Nevertheless, some accessions had %Ndfa values that were significantly larger than zero. These significant fixer accessions were of wild, landrace, and improved origin and derived up to 70% of their N from the atmosphere in 2023 and up to 45% in 2024 (Fig. 3). Accessions PI 628385 (an Olotón landrace from Chiapas, Mexico) and Ames 21890 (a non-crop progenitor, *Z. diploperennis*) were significant fixers in both years (Table 1; Fig. 3). No significant effect of accession was observed for N fixed (kg/ha – Fig S9).

**Figure 3.**
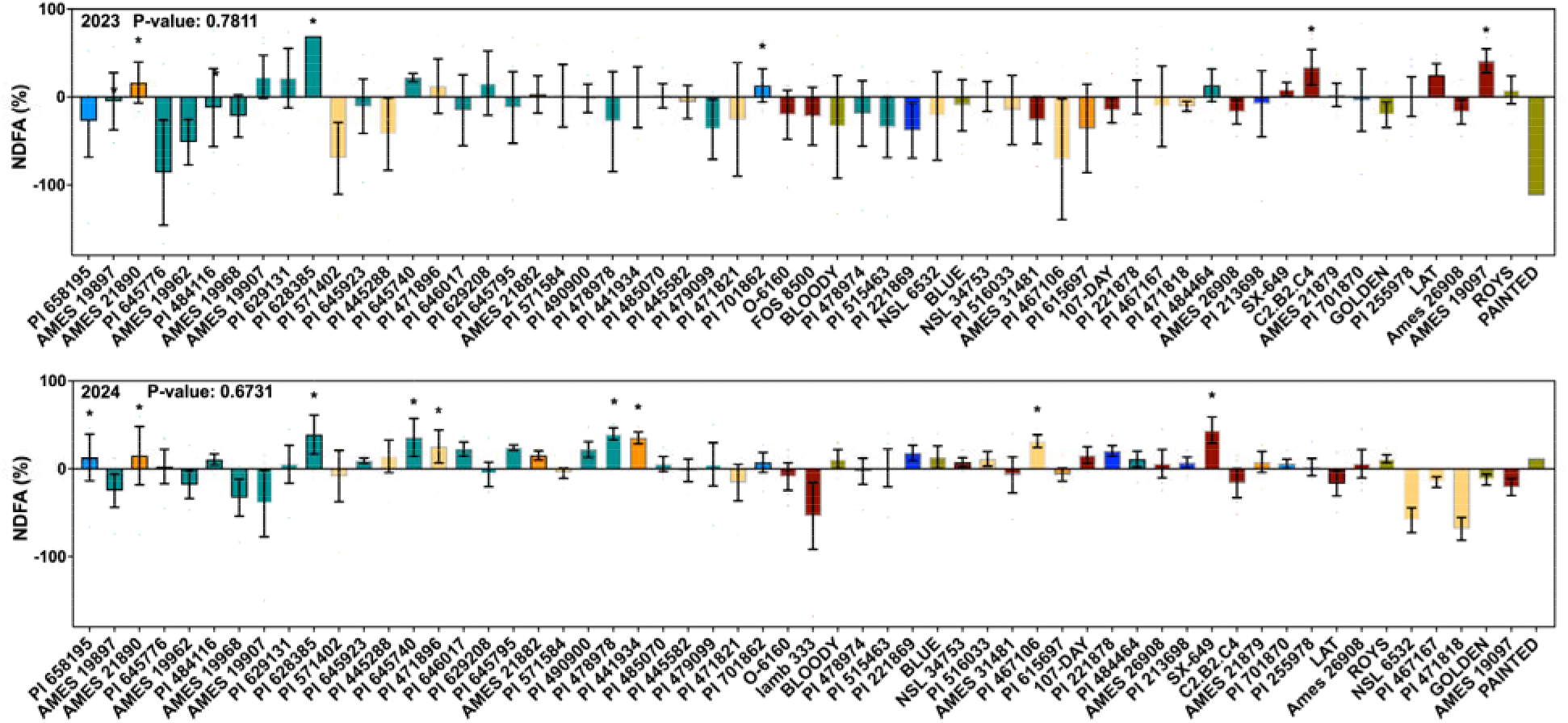
Variation in nitrogen fixation rate (using a natural abundance method) across diverse *Zea* accessions and domestication gradient. Accessions are ordered by ARLN in 2023 from Figure 2 and color-coded by domestication group. Least squares means estimates (with s.e. bars) were derived using generalized linear mixed models (GLMMs). Asterisks indicate the mean is significantly different than zero. Means that share a letter are not significantly different (Tukey’s adjustment, P < 0.001). Negative %Ndfa values indicate that the reference plants had lower δ^1^□N values than the focal accessions, suggesting that no biological nitrogen fixation occurred.

**Table 1.**
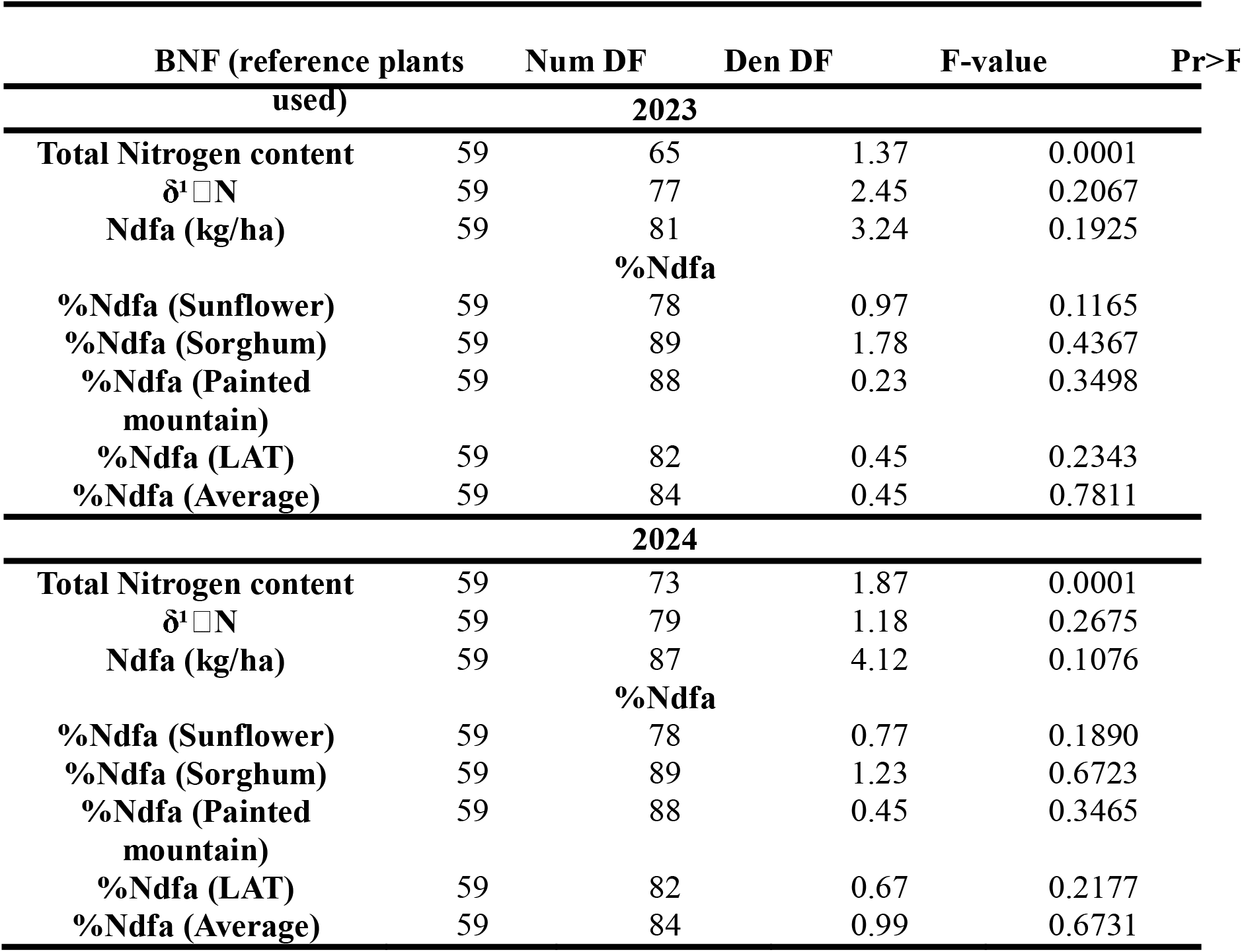
The effect of accession on N content and biological N fixation in 2023 and 2024.

Moreover, in both years, the majority of accessions did not differ significantly from the Midwest control for δ^15^N, %Ndfa, or nitrogen fixed (kg ha ¹) (Figs. S10–S12). For δ^15^N, three accessions NSL 6532 (a landrace from Paraguay), Lamb 333 (an improved variety thought to be a non-fixing control), and PI 471818 (a landrace from Bolivia)] differed significantly from the Midwest reference in 2024 (Fig. S10). Because the Dunnett contrasts were calculated as the Midwest reference minus each accession (Control − accession), the negative comparisons for these accessions indicate that they had significantly higher, rather than lower, δ^15^N values than the reference (Fig. S10). By contrast, no significant differences among accessions were detected for %Ndfa in either growing season (Fig. S11). Nevertheless, several accessions exhibited relatively large differences from the Midwest reference that approached statistical significance, particularly for %Ndfa in 2024. These trends included multiple landraces with an average difference of 23% (Dunnett’s P-values from 0.087 to 0.115), e.g., three landraces from Oaxaca, Mexico (Ames 19968, a Bolita landrace; and Ames 19907 and 19897, both Mixeño landraces), a Palomera Toluqueña landrace from Mexico, Mexico (Ames 19962), a Tabloncillo landrace from Jalisco, Mexico (PI 490900), three South American landraces (PI 471896 and PI 471818 from Bolivia and NSL 6532 from Paraguay), the improved line Ames 19097 (B73 × Mo17), and the heirloom variety Golden Bantam. For estimated nitrogen fixed (kg ha□¹), only a single accession differed significantly from the Midwest reference in 2024, showing significant higher estimated nitrogen fixed (kg ha□¹) (PI 490900, a Tabloncillo landrace from Jalisco, Mexico; Fig. S12).

### Domestication, spread, and improvement affect nitrogen-fixing traits

Our analysis revealed significant effects of the domestication gradient on several phenotypic traits of *Zea* taxa across two consecutive years (2023 and 2024), including number of nodes with aerial root (ARLN), plant height and biomass, and stem diameter (Table 2).

**Table 2.** The effect of domestication gradient (wild, landrace, heirloom, improved) on phenotypic traits of *Zea* taxa in 2023 and 2024.

| Trait | Num DF | Den DF | F-value | Pr>F |
| --- | --- | --- | --- | --- |
| 2023 |  |  |  |  |
| ARLN | 6 | 54.72 | 5.14 | 0.0002 |
| Aerial root diameter | 6 | 37.18 | 0.88 | 0.5279 |
| Mucilage | 6 | 48.56 | 3.46 | <0.0001 |
| Plant height | 6 | 53.43 | 7.54 | <0.0001 |
| First internode | 6 | 54.86 | 3.84 | 0.1800 |
| Second internode | 6 | 57.92 | 3.72 | 0.0022 |
| Stem diameter | 6 | 50.57 | 4.88 | 0.0003 |
| Plant biomass | 6 | 67.23 | 2.15 | <0.0001 |
| 2024 |  |  |  |  |
| ARLN | 6 | 55.41 | 8.50 | <0.0001 |
| Aerial root diameter | 6 | 51.69 | 1.37 | 0.2400 |
| Mucilage | 6 | 51.23 | 2.78 | <0.0001 |
| Plant height | 6 | 53.70 | 6.75 | <0.0001 |
| First internode | 6 | 52.87 | 4.31 | 0.0008 |
| Second internode | 6 | 53.63 | 6.36 | <0.0001 |
| Stem diameter | 6 | 50.74 | 7.46 | <0.0001 |
| Plant biomass | 6 | 62.78 | 4.11 | <0.0001 |

We observed a clear domestication gradient in the number of nodes bearing aerial roots, with ARLN declining with domestication and improvement (Table 2, Fig 4a). In 2023, the progenitors of cultivated maize produced the most aerial roots, followed by non-progenitor wilds and Mexican and South American landraces. In 2024, the difference between wild teosinte and landraces was not significant. In both years, heirloom and improved varieties exhibited significantly fewer nodes with aerial roots than teosinte, indicating that the domestication and improvement process has led to a reduction in this trait (Fig. 4a). By contrast, aerial root diameter did not change over the course of crop evolution (Table 2, Fig. 4b).

**Figure 4.**
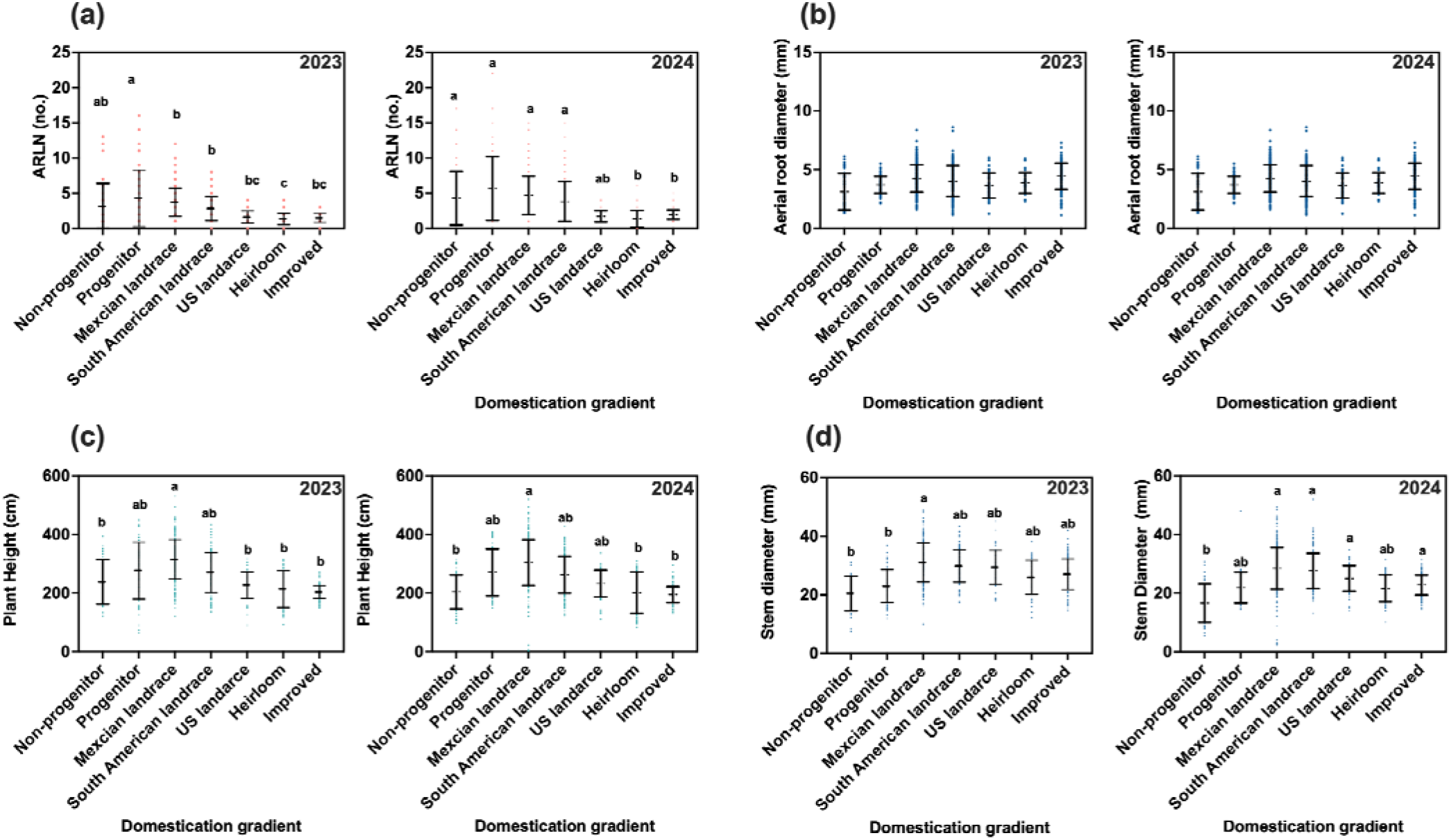
Variation in phenotypic traits associated with nitrogen fixation across the *Zea* domestication gradient in 2023 and 2024. Panels show least squares means (± SE) and individual domestication values for: (a) Number of nodes with aerial roots (ARLN), (b) Aerial root diameter (mm), (c) Plant height (cm), and (d) Stem diameter (mm). Categories represent stages along the domestication gradient: non-progenitor taxa, progenitors, Mexican landraces, South American landraces, U.S. landraces, heirloom varieties, and improved lines. Means that share a letter are not significantly different (Tukey’s adjustment, P < 0.001)

Mucilage production varied significantly across the domestication gradient, although the pattern differed between years (Fig. 5; Table 2). In 2023, production was generally low across all groups (ordinal scale 0–3), with non-progenitors and U.S. landraces exhibiting the lowest mucilage scores and wild progenitors and South American landraces showing higher values. By contrast, in 2024, Mexican landraces exhibited the highest mucilage production, non-progenitors the lowest, and all other groups were intermediate (Fig. 5). Thus, the effects of domestication, geographic spread, and improvement on mucilage production were inconsistent across years.

**Fig 5.**
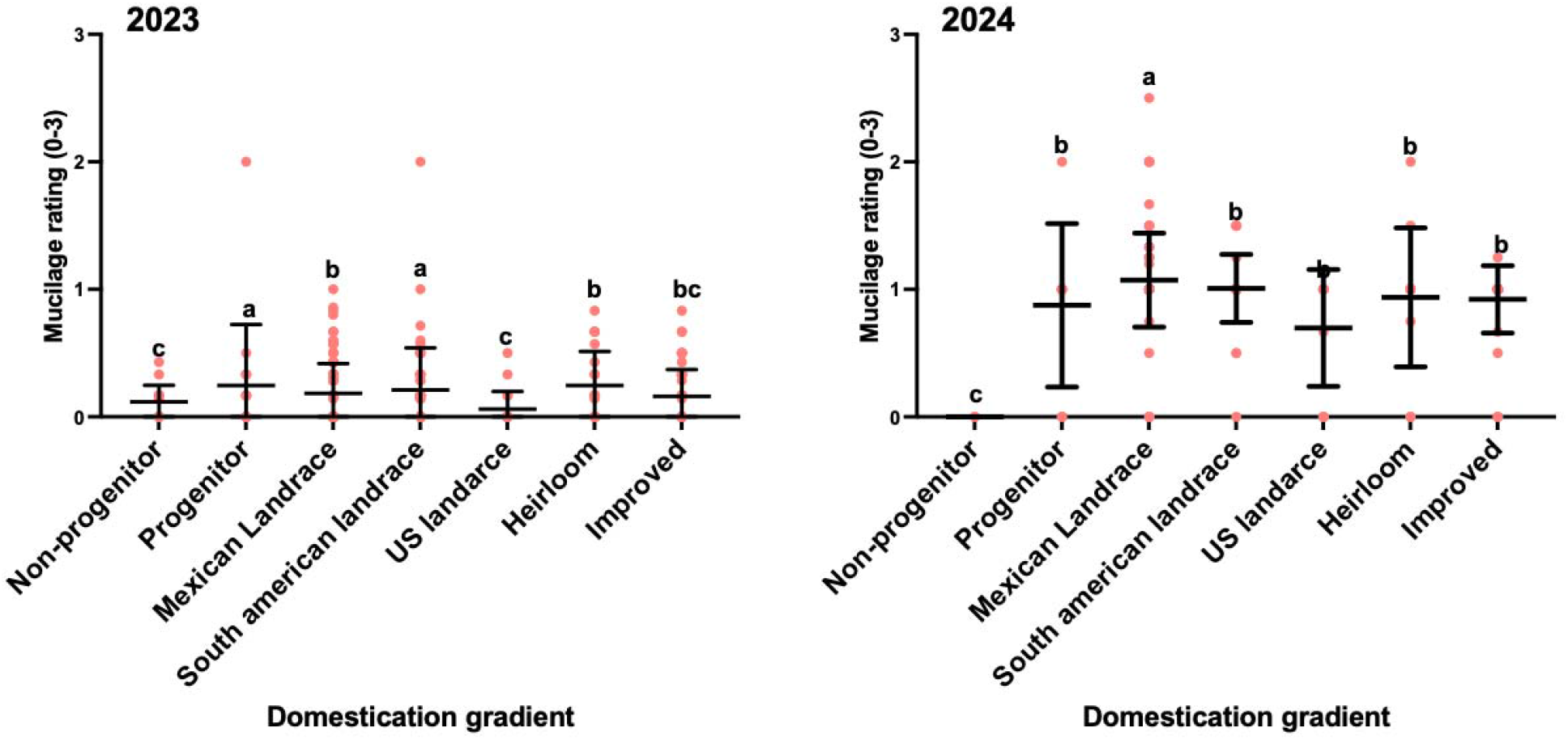
Variation in mucilage production across the domestication gradient of *Zea* taxa in 2023 and 2024. Mucilage production was scored on an ordinal scale from 0 to 3. Categories represent stages along the domestication gradient: non-progenitor taxa, progenitors, Mexican landraces, South American landraces, U.S. landraces, heirloom varieties, and improved lines. Means that share a letter are not significantly different (Tukey’s adjustment, P < 0.001).

The various stages of crop evolution have had clear impacts on measures of plant size. Mexican landraces consistently had the tallest plants across both years, but did not differ significantly from crop progenitors or other landraces, being only significantly taller than heirloom and improved varieties (Table 2; Fig. 4c). Thus, improvement seems to have shortened plants more than domestication and spread. By contrast, for stem diameter, while Mexican landraces exhibited the thickest stems in both years, this difference was only significant when compared to the thinner stemmed teosintes (both progenitor and non-progenitor in 2023, and only non-progenitor in 2024; Fig. 4d). Thus, domestication increased stem diameter but subsequent spread and improvement were less impactful. Interestingly, crop evolution appears to have progressively decreased plant biomass. In both years, wild teosintes (both progenitor and non-progenitor) had the highest amount of plant biomass and Mexican landraces had less biomass than wilds, indicating a domestication effect (Table 2; Fig S7); landraces from South America and the US had even less biomass that those form Mexico, indicating that geographic spread reduced biomass further. Finally, heirloom and improved maize had even less biomass, so improvement has also reduced this trait.

Internode traits exhibited year-dependent responses across the domestication gradient. First internode length differed among groups only in 2024, whereas second internode length differed in both years (Table 2; Fig. S13). Pairwise differences were inconsistent across years: U.S. landraces had shorter second internodes than several groups in 2023, whereas non-progenitors generally had shorter internodes than several landrace and heirloom groups in 2024. Overall, internode length showed no consistent directional pattern across the domestication gradient.

Total N (%) content differed significantly across the domestication gradient (Table 3). In both years, Mexican landraces exhibited the highest N content, significantly exceeding wild teosintes, while other cultivated groups were not different from or lower than Mexican landraces (Fig 6). Thus, domestication may have increased N content but further spread and improvement had a less prominent effect. Neither %Ndfa, nor Ndfa (kg/ha) – all estimates of biological N fixation – showed significant effects of domestication in either year (Table 3; Fig 7). Thus, crop evolution does not appear to have affected nitrogen fixation.

**Figure 6.**
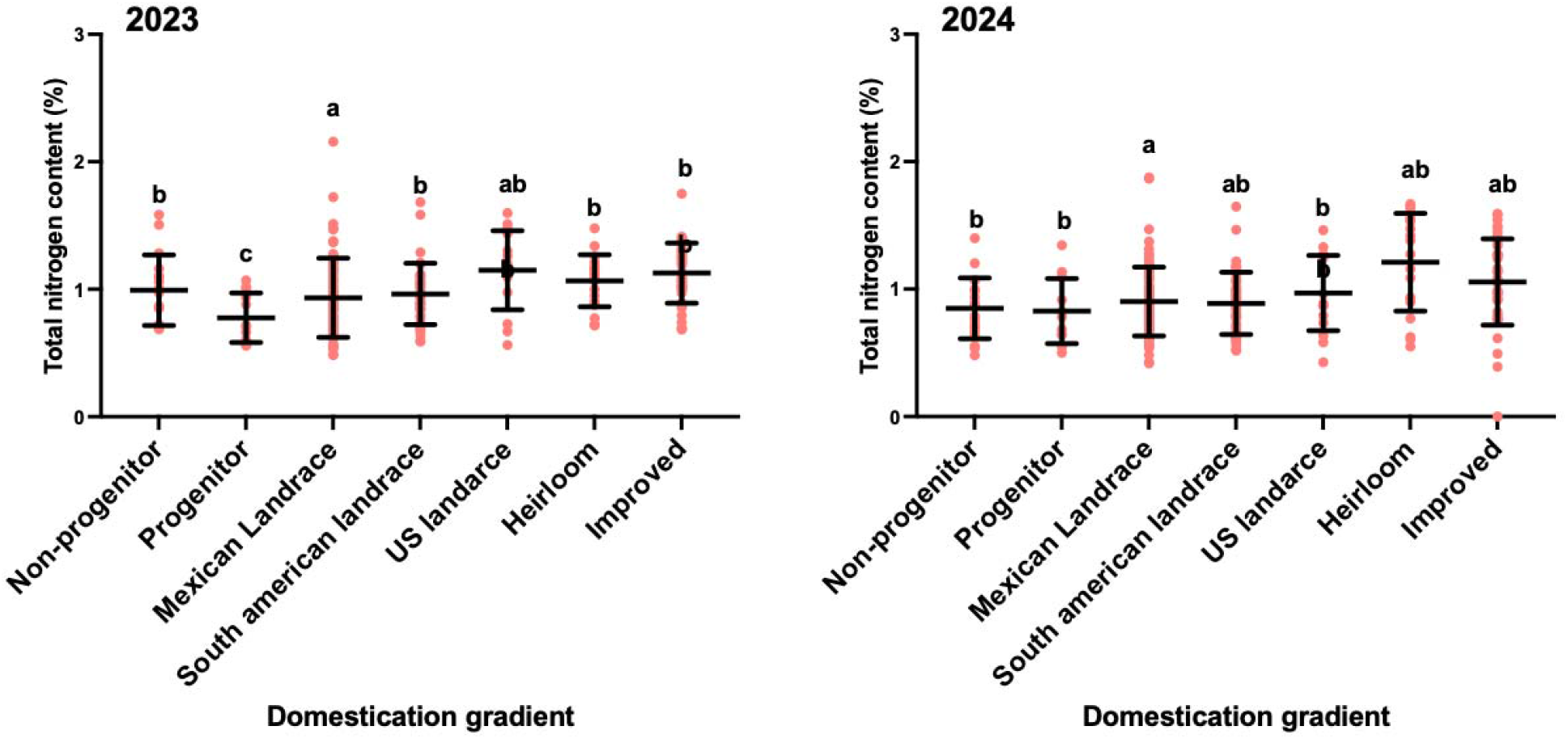
Variation in total nitrogen content (%) across the domestication gradient of *Zea* taxa in 2023 and 2024. Categories represent stages along the domestication gradient: non-progenitor taxa, progenitors, Mexican landraces, South American landraces, U.S. landraces, heirloom varieties, and improved lines. Letters above groups indicate significant differences among domestication categories based on post hoc multiple comparisons (P < 0.05).

**Figure 7.**
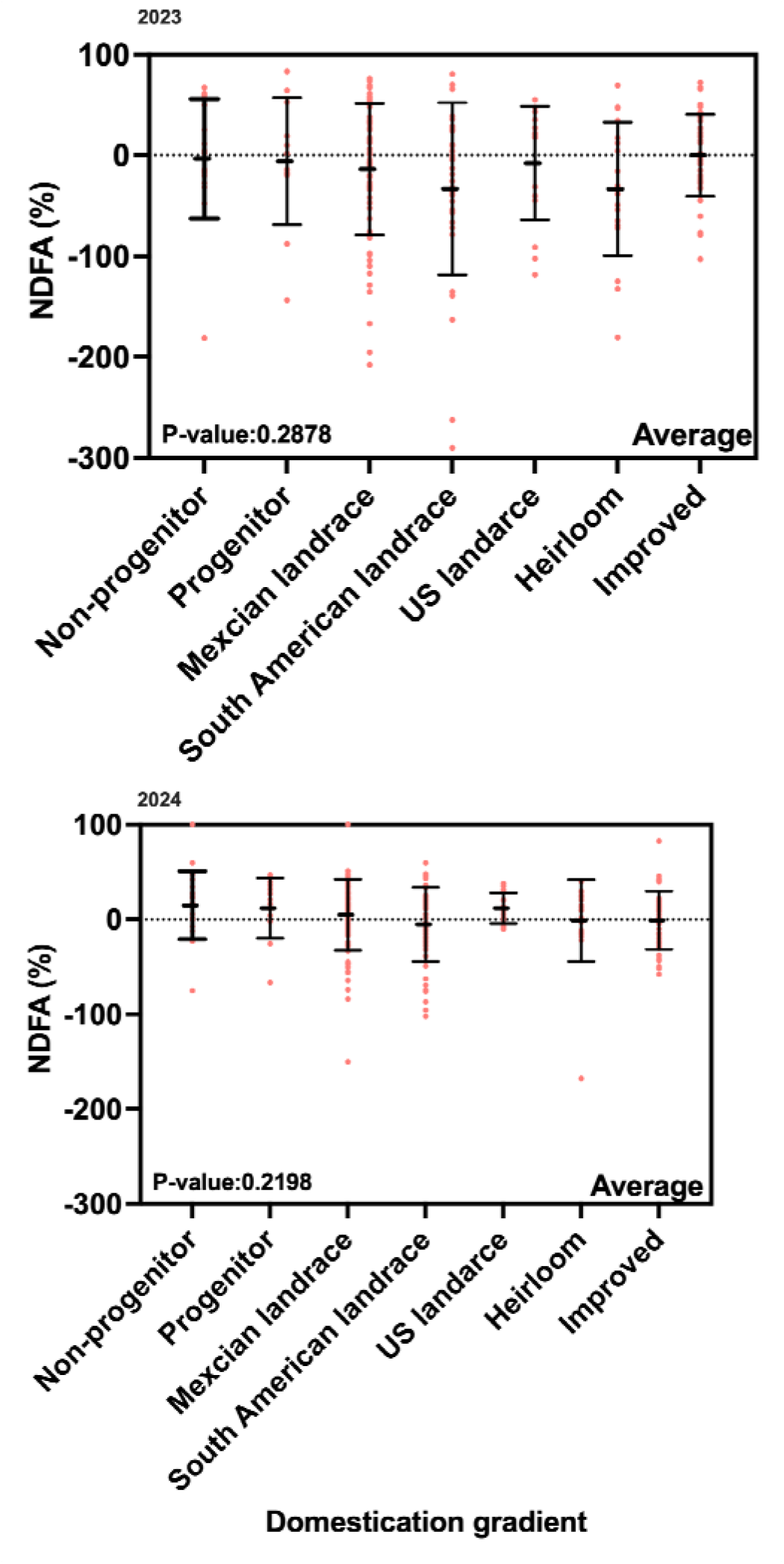
Variation in percent nitrogen derived from the atmosphere (Ndfa%) across the domestication gradient. P-values denote the effect of domestication gradient on percent nitrogen derived from the atmosphere (Ndfa%). Negative %Ndfa values indicate that the reference plants had lower δ^1^□N values than the maize plants, suggesting that no biological nitrogen fixation occurred.

**Table 3.**
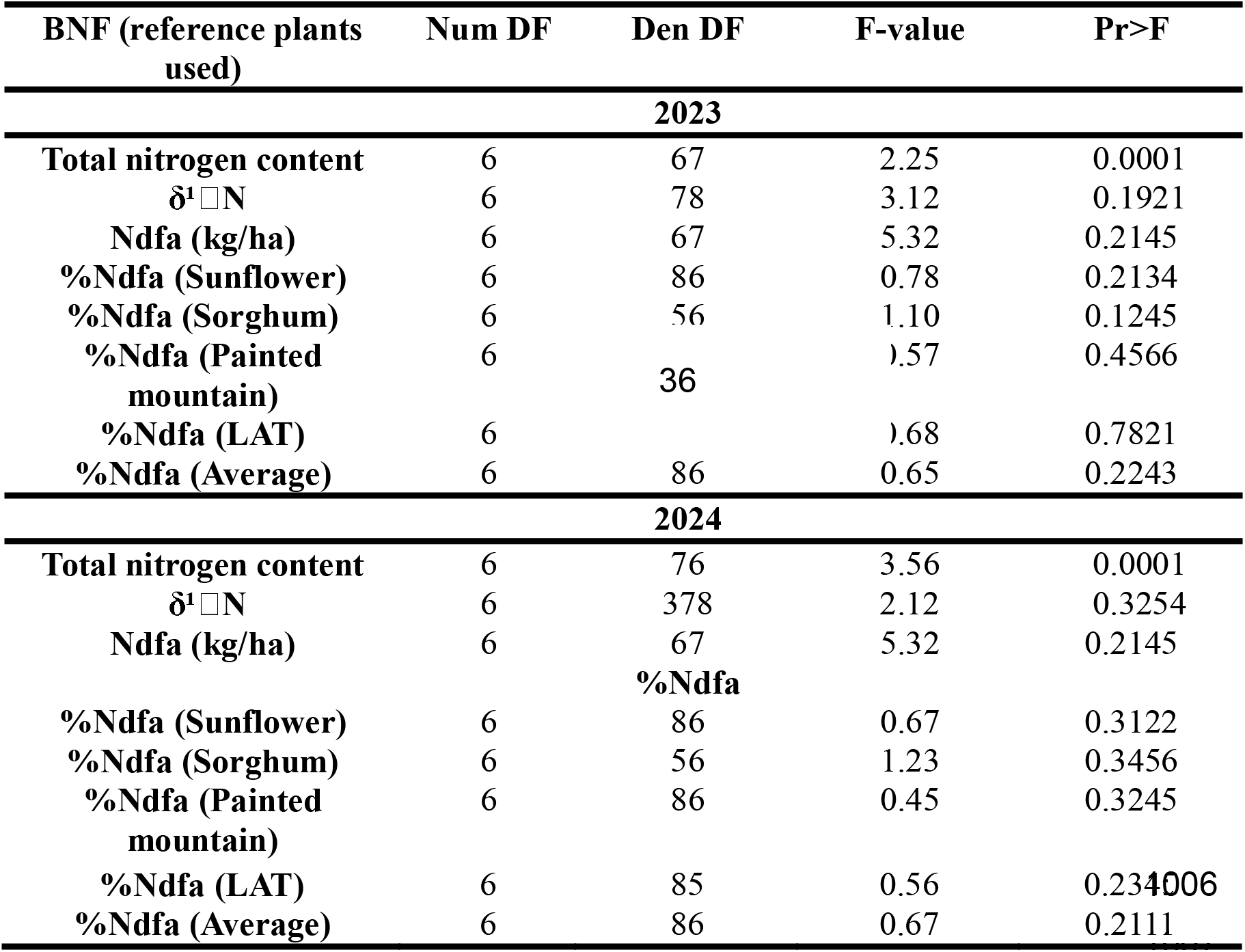
The effect of domestication on N content and biological N fixation in 2023 and 2024.

## Discussion

This study provides a multi-year, low-N, field evaluation of aerial root and growth traits and N derived from the atmosphere across diverse *Zea* accessions representing wild taxa (non-progenitor and progenitor); Mexican, South American, and U.S. landraces; heirloom varieties; and modern improved varieties. Accessions varied widely in aerial root phenotypes, but neither the proportion of N derived from the atmosphere (%Ndfa) nor the estimated amount of N derived from the atmosphere (Ndfa, kg ha ¹) differed significantly among accessions. Nevertheless, individual accessions from wild, landrace, and improved groups had showed significant N fixation in one or both years. Crop evolution appears to have reduced the number of nodes bearing aerial roots (ARLN) and aboveground biomass but has not altered %Ndfa nor Ndfa (kg/ha). Together, these results reveal a decoupling between the evolution of plant architecture and atmospheric N fixation: crop evolution substantially reshaped aerial-root and growth phenotypes without a corresponding directional change in N derived from the atmosphere. Thus, attempts to increase the capacity of maize to fix N will need to go beyond simple increases in aerial root production enhance the plant host, the microbial community, and the environmental conditions that can promote the association.

### Phenotypic diversity in N_2_ fixing traits among accessions

Substantial accession-level variation was evident in aerial root and mucilage traits. Wild *Zea* (progenitor and non-progenitors) and Mexican and South American landrace accessions had some of the highest numbers of nodes bearing aerial roots (Fig. 2). Mucilage production also varied significantly among accessions, although its expression differed between years, consistent with its environmental sensitivity (Pankievicz et al., 2022; Gao et al., 2023). This extensive within-group variation parallels the diversity previously documented in maize root architecture and host traits involved in plant–microbe interactions (López-Valdivia et al., 2022; Meier et al., 2022), suggesting that broad evolutionary categories capture only part of the phenotypic diversity relevant to potential diazotrophic associations.

Variation in these phenotypes, however, was not accompanied by significant accession-level differentiation in atmospheric N fixation. Neither δ^15^N nor %Ndfa differed significantly among accessions in either year, although individual accessions distributed across wild, landrace, and improved accessions had %Ndfa estimates significantly greater than zero (Fig. 3; Table 1). This broad distribution of detectable atmospheric N fixation is consistent with previous evidence for associative N fixation across Mexican landraces and other maize genotypes (Van Deynze et al., 2018; Goldstein & White, 2022; Gao et al., 2023; Connolly et al., 2025). Comparisons with maize varieties representative of U.S. Midwest production similarly provided little evidence for systematic differences; three accessions had lower δ^15^N than typical Midwest improved varieties in 2024, but no accession differed from the Midwest reference for %Ndfa; the one accession that did for Ndfa (kg ha□¹) in 2024 showed reduced Ndfa (Figs. S10–S12). Together, these results indicate that substantial phenotypic diversity in traits associated with diazotrophic niches does not necessarily translate into corresponding differentiation in whole-plant N derived from the atmosphere.

### Crop evolution reshaped aerial-root architecture but not atmospheric N fixation

Crop evolution strongly altered aerial-root architecture, but these changes occurred alongside broader shifts in plant form. The number of nodes bearing aerial roots (ARLN) declined from wild *Zea* and Mexican landraces toward heirloom and improved maize along with plant height and aboveground biomass which also declined across portions of this gradient (Fig. 4; Fig. S7). These patterns are consistent with the extensive remodeling of maize plant architecture during domestication and improvement, including changes in plant stature and root system architecture associated with cultivation and modern breeding (Duvick, 2005; Hufford et al., 2012; York et al., 2015; López-Valdivia et al., 2022; Ordóñez et al., 2024; Rinehart et al., 2024). Archaeological evidence further indicates that early maize retained more teosinte-like root systems (López-Valdivia et al., 2022), while comparisons among more recent germplasm likewise indicate evolutionary changes in nodal-root architecture (York et al., 2015; Schmidt et al., 2016; Guo et al., 2019; Rinehart et al., 2024). ARLN should therefore be considered within this broader evolutionary restructuring of maize architecture. Its decline could reflect direct selection on aerial-root development, but it could also have arisen indirectly through selection on correlated aspects of plant stature, phenology, or vegetative development.

Despite these pronounced changes in host phenotype, neither the proportion (%Ndfa) nor the estimated amount (Ndfa, kg ha ¹) of N derived from the atmosphere differed significantly among domestication categories in either year (Table 3; Fig. 7). Thus, evolutionary differentiation in aerial-root architecture was not accompanied by corresponding differentiation in atmospheric N fixation. This distinction is important because aerial roots and their mucilage can provide a carbon-rich, low-oxygen habitat for diazotrophs, and aerial-root characteristics have been associated with mucilage production, microbial colonization, and nitrogenase activity (Bennett et al., 2020; Pankievicz et al., 2022; Gao et al., 2023). Yet N fixation emerges from interactions among host phenotype, microbial communities, and environmental conditions rather than from aerial root architecture alone (Meier et al., 2022). Sierra Mixe maize demonstrates that atmospheric N fixation can make a substantial contribution to plant N (Van Deynze et al., 2018), while subsequent work has detected atmospheric N fixation across Mexican landraces that produce fewer aerial root bearing nodes than the extreme aerial root phenotype characteristic of Sierra Mixe maize (Connolly et al., 2025). Together with evidence from other maize accessions (Goldstein & White, 2022; Gao et al., 2023), these findings suggest that the ability to cultivate and benefit from atmospheric N fixation may be distributed across *Zea* diversity rather than following a directionally declining evolutionary trajectory from wilds to improved varieties.

More broadly, this decoupling illustrates why the evolution of associative N fixation cannot necessarily be inferred from the evolution of a single host trait. Domestication can alter root architecture, exudation, microbial recruitment, and other components of crop–microbiome interactions (Brisson et al., 2019; Udvardi et al., 2021; Meier et al., 2022; Chang et al., 2025), while maize evolution itself was not a linear transition from teosinte to modern cultivars. Geographic spread, admixture, introgression, farmer selection, and local adaptation generated substantial differentiation among maize populations, particularly among geographically separated landraces (Dominguez et al., 2024). Different components of the host–microbe association may therefore have followed partly independent evolutionary trajectories. Accordingly, the absence of differences in %Ndfa or Ndfa among our domestication categories should be interpreted as an absence of detectable differentiation under the environments evaluated here, rather than evidence that the biological mechanisms underlying associative N fixation remained unchanged throughout maize evolution.

### Decoupling of aerial root phenotypes and atmospheric N acquisition

Several mechanisms could explain why evolutionary differentiation in ARLN was not mirrored by differentiation in %Ndfa. First, the capacity of aerial roots to support diazotrophs depends on more than the number of aerial root-bearing nodes. Aerial root diameter, border-cell production, mucilage quantity and composition, and microbial colonization can influence the formation and activity of the diazotrophic microenvironment (Pankievicz et al., 2022; Gao et al., 2023). Genetic evidence further illustrates that these components of the aerial root–mucilage phenotype can be regulated independently. Gao et al. (2023) identified ZmSBT3, a subtilase that negatively regulates aerial-root mucilage secretion, and reported changes in alleles associated with increased mucilage secretion across teosinte and modern inbred germplasm. Domestication-related changes in root exudation, root traits, and microbiome-associated phenotypes have likewise been reported across other crop systems (Iannucci et al., 2017; Deng et al., 2024; He et al., 2025; Chang et al., 2025), consistent with broader evidence that domestication and breeding can modify traits underlying plant–microbe associations (Liu et al., 2020; Wen et al., 2021). Importantly, however, ZmSBT3 regulates mucilage secretion rather than ARLN itself and therefore should not be interpreted as a direct genetic explanation for the ARLN gradient observed here. Fewer aerial-root-bearing nodes in improved maize therefore need not translate proportionally into reduced atmospheric N fixation. Environmental conditions add another layer of variation by influencing the expression of traits that support diazotrophic associations. Mucilage production, for example, depends strongly on moisture and humidity (Van Deynze et al., 2018; Pankievicz et al., 2022), and its expression differed substantially between our two growing seasons. The composition and colonization of diazotrophic microbial communities may also vary among environments and locations, although we did not characterize microbial communities in this study. Consequently, differences in aerial root architecture may interact with both environmental conditions and locally available microbial communities to produce different realized diazotrophic associations across environments.

Second, atmospheric N fixation need not occur exclusively through the aerial root– mucilage system. The ^15^N natural-abundance approach estimates the contribution of atmospheric N at the whole-plant level but cannot identify the anatomical location or microbial pathway through which fixation occurred. Maize can associate with free-living, rhizosphere-associated, and endophytic diazotrophs in addition to those inhabiting aerial root mucilage (Goldstein & White, 2022; Dietrich et al., 2024). Contributions from these alternative association possibilities could therefore weaken the relationship between aerial root architecture and whole-plant %Ndfa. This is both a plausible explanation for the observed decoupling and an important limitation of our study; our isotopic measurements cannot partition atmospheric N generated from aerial root, rhizosphere, or endophytic associations. Combining isotopic tracing with compartment-specific nitrogenase measurements, microbial community characterization, and aerial and belowground root phenotyping would help resolve the relative contributions of these pathways.

### Implications for breeding and sustainability

These results suggest that breeding for associative N fixation will require a multi-trait approach rather than selection for a single visible phenotype. Aerial-root architecture is genetically variable in maize, and mucilage production and secretion also have a genetic basis, including regulation by genes such as ZmSBT3 (Gao et al., 2023). Host genetics can additionally influence root-associated microbial communities and plant–microbe interactions (Udvardi et al., 2021; Meier et al., 2022; Fonseca-García et al., 2024). Yet our results indicate that variation in these host traits should not be assumed to translate directly into variation in realized atmospheric N acquisition. Breeding strategies should therefore evaluate these components together—including aerial root architecture, mucilage production, microbial recruitment and activity, and reproducible atmospheric N acquisition—rather than selecting for aerial roots or mucilage alone. Plant and microbial components of these associations provide complementary targets for improvement (Wen et al., 2021; Gao et al., 2023; Fonseca-García et al., 2024; Chang et al., 2025). Identifying germplasm in which favorable host traits are coupled with reproducible atmospheric N acquisition across environments will ultimately be necessary to determine whether associative N fixation can contribute meaningfully to lower-input maize production.

Our study implies that traits potentially contributing to associative N fixation, including aerial root development and mucilage production, are present but unevenly distributed across maize germplasm. Domestication, geographic spread, and improvement have altered some of these traits, as has been observed for plant–diazotroph associations in other crops (e.g., Fernandes Júnior et al., 2013). Our inclusion of germplasm from a N-efficiency breeding program provided an informative comparison. C2.B2.C46, FOS 8500, Lamb 333, and LAT7-113 × LH123-13 were included as varieties representing contrasting N-use-efficiency characteristics (Goldstein et al., 2019; Goldstein, 2022; Goldstein & White, 2026). This variation among improved germplasm, together with the absence of a significant overall accession effect on %Ndfa, suggests that atmospheric N acquisition can vary even among improved varieties and emphasizes the need to validate accession-level signals across years and environments before identifying germplasm as reliable breeding targets. These observations are intriguing, although they should be interpreted cautiously: %Ndfa did not differ significantly among accessions overall, and C2.B2.C46 was identified as a significant fixer in both growing seasons. Nevertheless, the comparatively high %Ndfa signal of C2.B2.C46, together with the significant atmospheric N acquisition detected in SX-649 in 2024, illustrates variation in atmospheric N acquisition among improved germplasms. In contrast, a non-fixing improved control (Lamb 333) exhibited significantly higher δ^15^N than the Midwest reference in 2024, a pattern consistent with lower atmospheric N acquisition relative to those conventional controls. Together, these contrasting responses suggest that germplasm developed or evaluated for performance under N limitation does not necessarily exhibit consistently greater atmospheric N acquisition, but may provide useful material for further testing of relationships among N-use efficiency, plant–microbe interactions, and associative N fixation.

Recent genomic studies further indicate that many South American landraces remain underutilized in breeding despite harboring substantial genetic diversity and locally adapted alleles of potential agronomic value (Dominguez et al., 2024). The diversity represented by landraces and wild *Zea* therefore provides an important resource for evaluating and improving associative N fixation, but our results suggest that breeding targets should not be defined by any single component of this association. Rather than selecting specifically for greater aerial root production, mucilage production, or %Ndfa, breeding programs could evaluate combinations of host traits, microbial associations, and reproducible atmospheric N fixation across environments. Germplasm such as Sierra Mixe demonstrates the potential for substantial atmospheric N acquisition in maize (Van Deynze et al., 2018), while the broader variation represented in diverse maize germplasm provides opportunities to determine which combinations of plant and microbial traits produce stable benefits under N-limited conditions. Incorporating this diversity into breeding populations, particularly within regions where locally adapted landraces are already cultivated, could help identify genetic backgrounds in which associative N fixation contributes to N acquisition without compromising agronomic performance. Such an approach places germplasm diversity, rather than any single phenotype or specific engineering strategy, at the center of efforts to evaluate associative N fixation as a potential component of lower-input maize production.

## Conclusion

This study provides one of the first systematic field evaluations of traits associated with biological N fixation across a broad representation of the maize domestication, geographic spread, and improvement gradient. Crop evolution was associated with substantial changes in plant phenotype, most notably a reduction in the number of nodes bearing aerial roots (ARLN), whereas mucilage production showed year-dependent patterns. In contrast, neither the proportion (%Ndfa) nor the estimated amount (Ndfa, kg ha ¹) of N derived from the atmosphere differed significantly among domestication groups. At the accession level, %Ndfa likewise did not differ significantly among accessions, although individual accessions from wild, landrace, and improved germplasm exhibited detectable atmospheric N acquisition. Together, these results reveal that evolutionary changes in aerial root architecture were not accompanied by a corresponding directional change in whole-plant atmospheric N fixation. More broadly, our findings emphasize that associative N fixation should not be inferred from any single host phenotype. Aerial root architecture, mucilage production, microbial associations, environmental conditions, and atmospheric N fixation may represent interacting but partly independent components of the phenotype. The diversity observed across *Zea* germplasm therefore provides material for further testing rather than evidence that high BNF capacity has been retained or lost during crop evolution. Future work should evaluate promising germplasm across environments and reduced-N conditions, determine the relative contributions of aerial root, rhizosphere, and endophytic pathways to atmospheric N fixation, and identify combinations of host and microbial traits that produce reproducible N inputs. Establishing these relationships will be necessary before associative N fixation can be effectively incorporated into breeding or microbiome-based strategies aimed at reducing synthetic N inputs in maize production.

## Acknowledgements

We thank the staff of the Waterman Agricultural and Natural Resources Laboratory and members of the Mercer Lab, particularly Edison Cigany, Meena Jayaraman, Brittany Lewis, and Yudai Takenaka, for their assistance with fieldwork during the 2023 and 2024 growing seasons. We also thank the USDA North Central Regional Plant Introduction Station (NCRPIS) and the International Maize and Wheat Improvement Center (CIMMYT) for providing germplasm. This research was supported by the USDA National Institute of Food and Agriculture (grant no. 20226701337080). The funder had no role in the study design; data collection, analysis, or interpretation; manuscript preparation; or the decision to submit the manuscript for publication.

## Author contributions

KeM performed the experiments and collected the data, analysed the data, and drafted the manuscript. LC, NL and TA performed the experiments and collected the data and revised the manuscript. KM, and RD funding acquisition and supervision. KM, RD, AL, VB, and HP revised the manuscript for important intellectual content and discussed the core idea of the project.

## Data availability

Data will be made publicly available upon acceptance of this manuscript.

