## Supplementary file for "Domestication, spread, and improvement have shaped biological nitrogen fixation traits in *Zea* taxa"

**Trait measurement**

The traits used in this study were measured using standardized methods tailored to each trait. To obtain trait data, three plants per subplot marked for harvest were cut at the R1–R2 growth stages (silking to blister). At this time, plants had fully developed aerial roots and mucilage secretion passed its peak, allowing consistent measurements across accessions.

***Plant height**:* This trait was measured from the base of the plant to the collar of the newest fully formed leaf. Data collection for plant height was conducted at the time of harvest (the R1–R2 growth stages).

***Number of nodes with aerial roots:*** The quantification of visible nodal roots and nodes on each plant was conducted manually. This study employed the numbering system of aerial root layer number (ARLN) as described by Zhang *et al.* (2018). In this system, the base layer of brace roots at the soil surface was designated as '0'. Subsequent layers above the soil surface, including the second brace root layer and any aerial nodes, was numbered starting from '2' and upwards. This methodical approach ensures a standardized and replicable measurement of nodal roots and nodes across all plants (the R1–R2 growth stages).

***Diameter of Aerial roots:*** The thickness of nodal roots was measured using calipers. Root diameter measurements was taken on three different roots, utilizing a degree-based system to standardize the process. In this system, we conceptualize the arrangement of roots around the stem as if they were positioned along the degrees of a circle. Measurements were conducted at specific positions of 0°, 120°, and 240°. Calipers were placed approximately 2 cm below the point where each root emerged from the main stem. By selecting roots at different angular positions around the stem, we ensured a representative sampling of root orientations. Whenever possible, the youngest aerial roots emerging from higher nodes were measured.

***Length of the first and second internodes:*** The lengths of the internodes was measured Specifically, the first internode length was determined by measuring the distance between the first two nodes with aerial roots, while the second internode length was assessed by measuring the distance between the second and third nodes with aerial roots. This measurement was taken from the base of one node to the top of the subsequent node, ensuring accuracy and consistency in the data collected.

***Mucilage rating****:* This trait was evaluated through a qualitative assessment using a predefined scale designed to capture both the amount and consistency of mucilage produced by each plant. Mucilage production was rated on a scale from 0 to 3, where a rating of 0 indicates absence of mucilage and 3 signifies abundant mucilage visibly dripping from aerial roots (Fig. S2). Observations were conducted once per week throughout the growing season and additionally following rainfall events, as increased relative humidity promotes mucilage secretion. Assessments were focused on younger aerial roots, which are more likely to exude mucilage than older nodes. The resulting mucilage ratings were subsequently used as input for logistic regression analyses to model the probability of mucilage occurrence across accessions and environments.

### Nitrogen fixation measurement

#### ***$^{15}N$ Natural Abundance***

The ratios of nitrogen isotopes in different maize accessions were analyzed using whole plant material. The $^{15}N$ natural abundance method relies on the relative abundance of the stable isotope $^{15}N$ in the atmosphere and soil, with $^{15}N$ abundance being more abundant in the soil than in the air. As a consequence, plants that derive N from the atmosphere will exhibit reduced $\delta^{15}N$ levels when compared to reference non-N-fixing plants. Plants were harvested at the VT and R1 (before flowering) stages and processed through a woodchipper specifically designated for this purpose. The wet weight of the plant biomass was recorded before drying the samples completely at 55°C, after which the dry weight was noted. After chippin, three whole plants (where available), a subsample of the homogenized tissue was taken and placed in a dryer until the samples reach a consistent level of dryness. These dry subsamples were ground using a Udy Mill grinder, optimized for the rapid, uniform grinding of a wide variety of plant samples, including conventional corn and sunflower, which are included for comparative purposes. The grinder was meticulously cleaned between uses with a vacuum, air compressor, and 80% ethanol to minimize cross-contamination between samples from different subplots. Each sample was ground to a fine, 100-mesh powder, and approximately 10-20g of each sample was processed to this degree of fineness.

For the $\delta^{15}N$ analysis, we employed the natural abundance method to measure the extent of atmospheric $N_2$ fixation in the plants. This method was based on the ratio of naturally occurring $\delta\ ^{15}N$ and $\delta\ ^{14}N$ isotopes. By comparing the $\delta^{15}N$ N ratio of a focal accession with reference plants, we can calculate the percentage of nitrogen derived from the atmosphere (%Ndfa = [($\delta^{15}N$ reference - $\delta^{15}N$ focal)/ ($\delta^{15}N$ reference – B)] x 100), where B represents the $\delta^{15}N$ abundance in air, assumed to be zero. For the isotopic analysis, approximately 5 mg of plant tissue was weighed into tin capsules and combusted in an elemental analyzer (CHN EA 1108, Carlo Erba Instruments, Italy) at temperatures up to 1800°C. The resulting nitrogen oxides (NOx) were converted to nitrogen ($N_2$) and then separated using a chromatographic column. A portion of the $N_2$ gaswas directed to a thermal conductivity detector for % $N_2$ quantification, while another portion was analyzed for $\delta^{15}N$ isotopic composition using a Delta V Advantage Isotope Ratio Mass Spectrometer (IRMS). The $\delta^{15}N$ abundances were expressed as $\delta^{15}N$ [‰], calculated as [(R_sample - R_standard) - 1] * 1000, where R represents the $\delta^{15}N$ / $\delta\ ^{14}N$ ratio (atom %) of the sample and the standard. The standard for this measurement is atmospheric $\delta^{15}N_2$ , with an $\delta^{15}N$ abundance of 0.00366295. This detailed and systematic approach to $\delta^{15}N$ isotope measurements enablde us to accurately assess the nitrogen fixation component of the different Zea taxa.

**Table S1.** List of accessions used in this study.

| Accession ID | Name | Domestication type | Taxon | Elevation (m) | Origin |
|---|---|---|---|---|---|
| PI 658195 | - | Teosinte | Zea mays subsp. mexicana | 2425 | México, Mexico |
| PI 701870 | - | Teosinte | Zea mays subsp. parviglumis | 646 | México, Mexico |
| PI 615697 | - | Teosinte | Zea nicaraguensis | 9 | Chinandega, Nicaragua |
| Ames 21879 | - | Teosinte | Zea luxurians | 800 | Chiquimula, Guatemala |
| PI 441934 | - | Teosinte | Zea mays subsp. huehuetenangensis | 1350 | Huehuetenango, Guatemala |
| Ames 21890 | - | Teosinte | Zea diploperennis | 1950 | Jalisco, Mexico |
| Ames 21882 | - | Teosinte | Zea perennis | 2150 | Jalisco, Mexico |
| PI 701862 | - | Teosinte | Zea mays subsp. mays subsp. mexicana | 540 | México, Mexico |
| PI 645740 | Comiteco | Mexican Landrace | - | 1840 | Chiapas, Mexico |
| Ames 19968 | Bolita | Mexican Landrace | - | 1400 | Oaxaca, Mexico |
| PI 478974 | Tepecintle | Mexican Landrace | - | 475 | Chiapas, Mexico |
| PI 478978 | Zapalote Chico | Mexican Landrace | - | 50 | Chiapas, Mexico |
| Ames 19897 | Mixeno | Mexican Landrace | - | 1700 | Oaxaca, Mexico |
| Ames 19907 | Mixteco | Mexican Landrace | - | 2500 | Oaxaca, Mexico |
| PI 645776 | Ancho | Mexican Landrace | - | 1400 | Guerrero, Mexico |
| PI 484116 | Cacahuazintle | Mexican Landrace | - | 2500 | Michoacán de Ocampo, Mexico |
| PI 485070 | Chalqueno | Mexican Landrace | - | 2229 | Puebla, Mexico |
| Ames 19962 | Palomero Toluqueno | Mexican Landrace | - | 2100 | México, Mexico |
| PI 629208 | Conico | Mexican Landrace | - | 2240 | México, Mexico |
| PI 490900 | Tabloncillo | Mexican Landrace | - | 870 | Jalisco, Mexico |
| PI 479099 | Nal-Tel | Mexican Landrace | - | 30 | Yucatán, Mexico |
| PI 645795 | Celaya | Mexican Landrace | - | 1760 | Guanajuato, Mexico |
| PI 645923 | Jala | Mexican Landrace | - | 1097 | Nayarit, Mexico |
| PI 646017 | Tuxpeno Norteno | Mexican Landrace | - | 213 | Tamaulipas, Mexico |
| PI 484464 | Cristalino de Chihuahua | Mexican Landrace | - | 2411 | Chihuahua, Mexico |

| | | | | | |
|---|---|---|---|---|---|
| PI 628385 | Oloton | Mexican Landrace | - | 1870 | Chiapas, Mexico |
| PI 629131 | Tuxpeno | Mexican Landrace | - | 690 | Chiapas, Mexico |
| PI 515463 | Cacahuazintle | Mexican Landrace | - | 410 | Sonora, Mexico |
| PI 255978 | Rhode Island White Cap Flint | US Landrace | - | - | Rhode Island, United States |
| PI 221869 | Johnson County White | US Landrace | - | 450 | Missouri, United States |
| PI 221878 | Leaming | US Landrace | - | 93 | Missouri, United States |
| PI 213698 | Reids Yellow Dent | US Landrace | - | - | Indiana, United States |
| PI 467167 | CHZM 05 017 | South American Landrace | - | 250 | Valparaíso, Chile |
| PI 471821 | Madre de Dios 22 | South American Landrace | - | 500 | Brazil |
| PI 471818 | Bolivia 47 | South American Landrace | - | 362 | Bolivia |
| NSL 6532 | Paraguay 136 | South American Landrace | - | 366 | Caaguazú, Paraguay |
| PI 445582 | Valle 347 | South American Landrace | - | 1300 | Valle de Cauca, Colombia |
| PI 516033 | Caldas 367 | South American Landrace | - | 1540 | Argentina |
| PI 471896 | Bolivia 1048 | South American Landrace | - | 1948 | Bolivia |
| PI 445288 | Putumayo 345 | South American Landrace | - | 2000 | Putumayo, Colombia |
| PI 571402 | Ancash 285 | South American Landrace | - | 2300 | Ancash, Peru |
| PI 571584 | Apurimac 83 | South American Landrace | - | 2900 | Apurímac, Peru |
| PI 467106 | Cuzco 18 | South American Landrace | - | 1450 | CHL, Chile |
| Ames 19097 (B73 x Mo17) | - | Improved | - | - | - |
| Ames 31481 (PHB47 x PHZ51) | - | Improved | - | - | - |
| Ames 26908 (US13) | - | Improved | - | - | - |
| NSL 34753 (Hickory king) | - | Improved | - | - | - |
| C2.B2.C46 | - | Improved – bred for NUE | - | - | - |

| | | | | | |
|---|---|---|---|---|---|
| FOS 8500 | - | Improved – non-fixing | - | - | - |
| Lamb333 | - | Improved – non-fixing | - | - | - |
| O-6160 | - | Improved | - | - | Ohio, United States |
| 107-DAY | - | Improved | - | - | Ohio, United States |
| SX-649 | - | Improved | - | - | - |
| LAT7-113 x LH123-13 (LAT) | - | Improved – non-fixing | - | - | - |
| Bloody Butcher (BLOODY) | - | Heirloom | - | - | - |
| Blue Clarage (BLUE) | - | Heirloom | - | - | - |
| Roy's Calais Flint Corn (ROYS) | - | Heirloom | - | - | - |
| Painted Mountain (PAINTED) | - | Heirloom - non-fixing | - | - | - |
| Golden Bantam (GOLDEN) | - | Heirloom | - | - | - |
| Sorghum Sudangrass (SS) | - | Reference | - | - | - |
| Mammoth Sunflower (MS) | - | Reference | - | - | - |

**Table S2.** The effect accession on phenotypic traits of *Zea* taxa in 2023. A generalized linear mixed models with accession being as a fixed effect and block as random effects were used to estimate the effect accession on phenotypic traits of *Zea* taxa.

| Trait | Num DF | Den DF | F-value | Pr>F |
|---|---|---|---|---|
| **2023** | | | | |
| **ARLN** | 59 | 667 | 10.75 | <0.0001 |
| **Aerial root diameter** | 59 | 441 | 4.82 | 0.7682 |
| **Mucilage** | 59 | 421 | 4.67 | <0.0001 |
| **Plant height** | 59 | 676 | 23.21 | <0.0001 |
| **Stem diameter** | 59 | 677 | 11.18 | <0.0001 |
| **First internode** | 59 | 662 | 6.41 | <0.0001 |
| **Second internode** | 59 | 662 | 4.08 | <0.0001 |

**Table S3.** The effect accession on phenotypic traits of *Zea* taxa in 2024. A generalized linear mixed models with accession being as a fixed effect and block as random effects were used to estimate the effect accession on phenotypic traits of *Zea* taxa.

| Trait | Num DF | Den DF | F-value | Pr>F |
|---|---|---|---|---|
| **2024** | | | | |
| **ARLN** | 59 | 837 | 16.36 | <0.0001 |
| **Aerial root diameter** | 59 | 678 | 0.58 | 0.9944 |
| **Mucilage** | 59 | 654 | 12.36 | <0.0001 |
| **Plant height** | 59 | 847 | 25.96 | <0.0001 |
| **Stem diameter** | 59 | 846 | 17.82 | <0.0001 |
| **First internode** | 59 | 837 | 5.99 | <0.0001 |
| **Second internode** | 59 | 837 | 8.78 | <0.0001 |

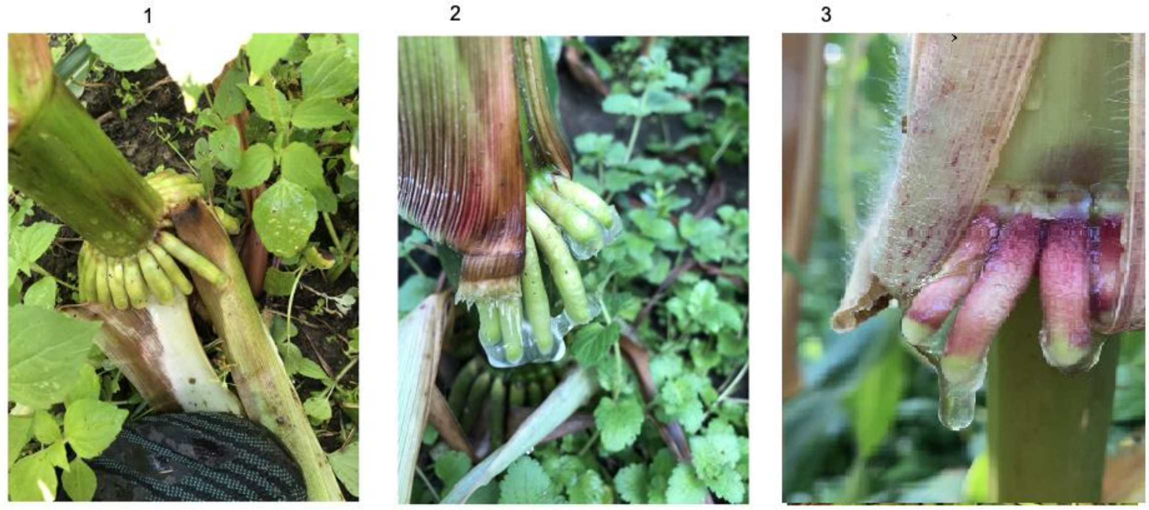


**Figure S1.** mucilage rating system used in this study. The photo shows an ordinal scale ranging from 1 to 3. The photos were taken by Keyvan Maleki.

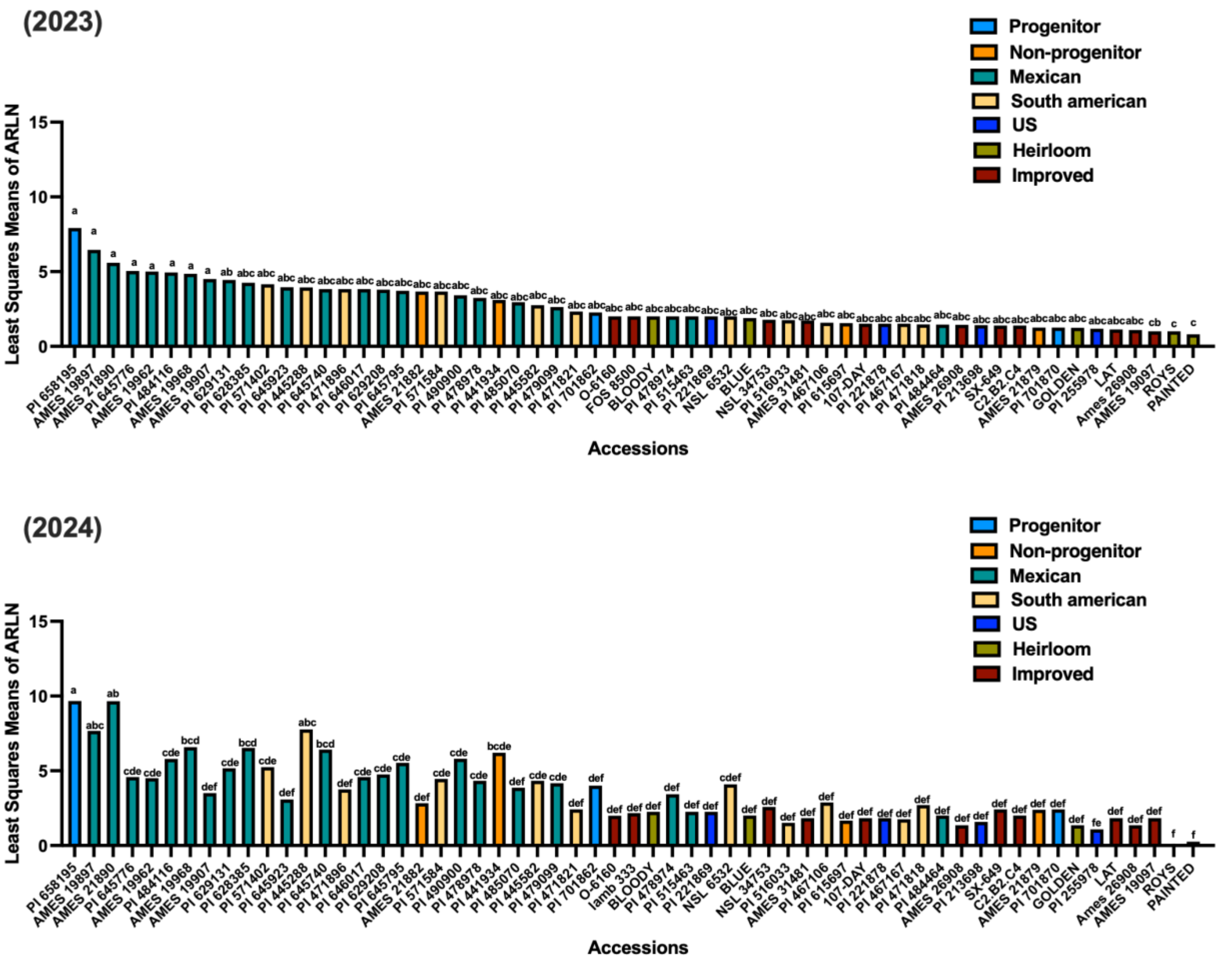


**Figure S2.** Least-squares means of aerial root node layers (ARLN) across maize accessions representing different domestication groups (Progenitor, Non-progenitor, Mexican landraces, South American landraces, U.S. landraces, Heirloom, and Improved lines). Bars are colored by domestication group and ordered by descending mean ARLN. The panels show accessions without grouping letters and Tukey–Kramer multiple comparison groupings ($\alpha = 0.05$); accessions sharing the same letter are not significantly different. Least-squares means were estimated from a generalized linear mixed model with a negative binomial distribution and log link, accounting for block as a random effect.

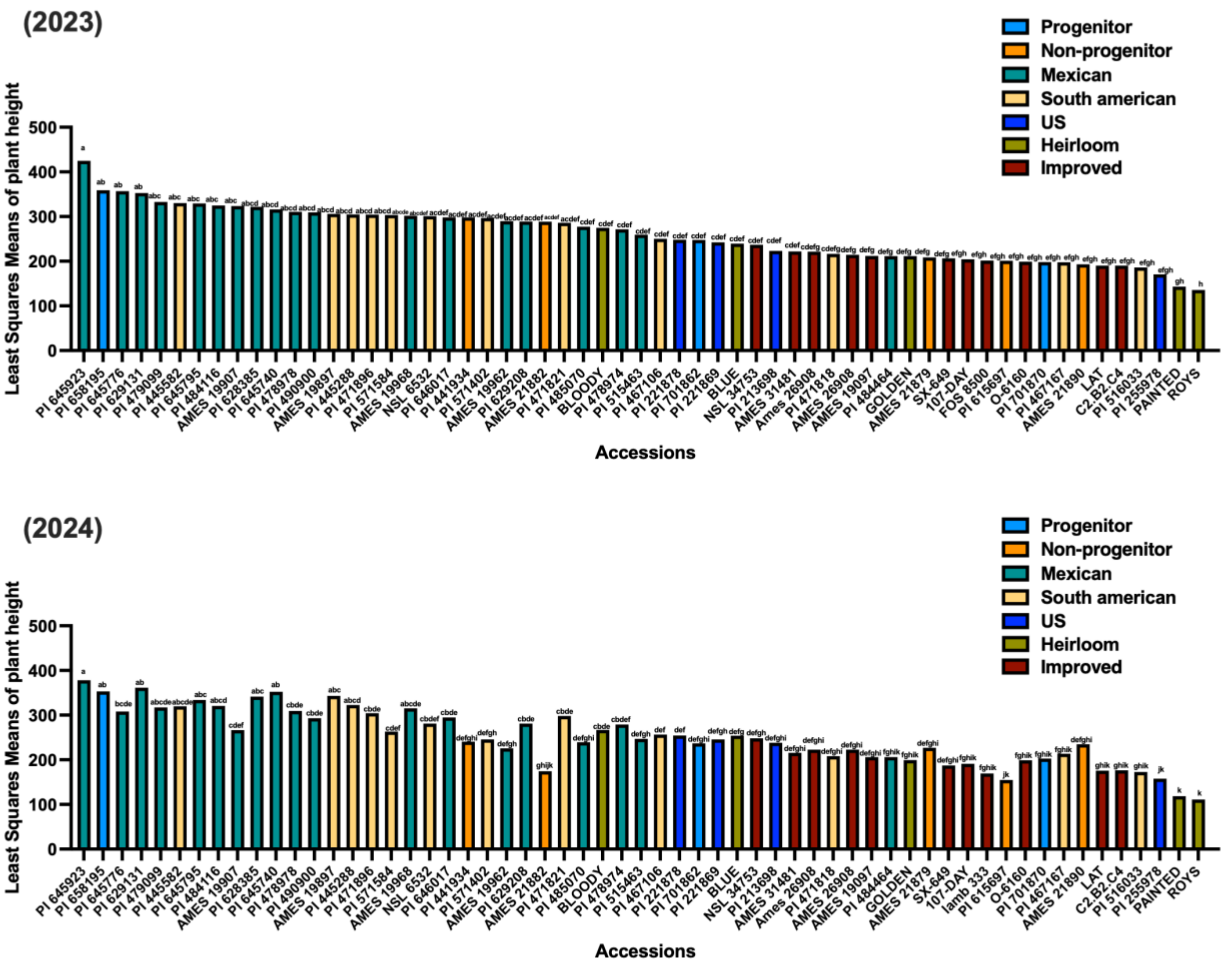


**Figure S3.** Least-squares means of plant height across maize accessions representing different domestication groups (Progenitor, Non-progenitor, Mexican landraces, South American landraces, U.S. landraces, Heirloom, and Improved lines). Bars are colored by domestication group and ordered by descending mean ARLN. The panels show accessions without grouping letters and Tukey–Kramer multiple comparison groupings ($\alpha = 0.05$); accessions sharing the same letter are not significantly different. Least-squares means were estimated from a generalized linear mixed model with a negative binomial distribution and log link, accounting for block as a random effect.

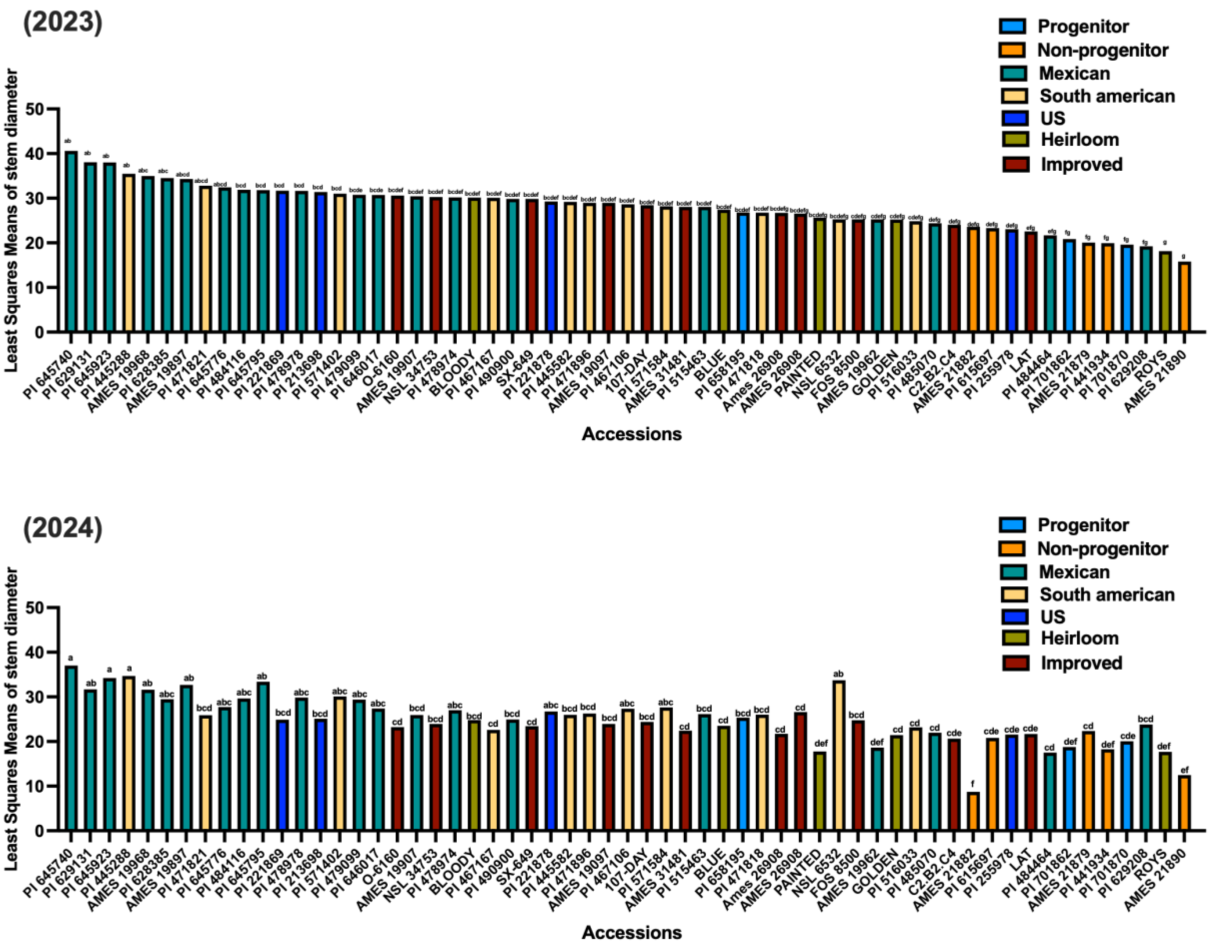


**Figure S4.** Least-squares means of stem diameter across maize accessions representing different domestication groups (Progenitor, Non-progenitor, Mexican landraces, South American landraces, U.S. landraces, Heirloom, and Improved lines). Bars are colored by domestication group and ordered by descending mean ARLN. The panels show accessions without grouping letters and Tukey–Kramer multiple comparison groupings (α = 0.05); accessions sharing the same letter are not significantly different. Least-squares means were estimated from a generalized linear mixed model with a negative binomial distribution and log link, accounting for block as a random effect.

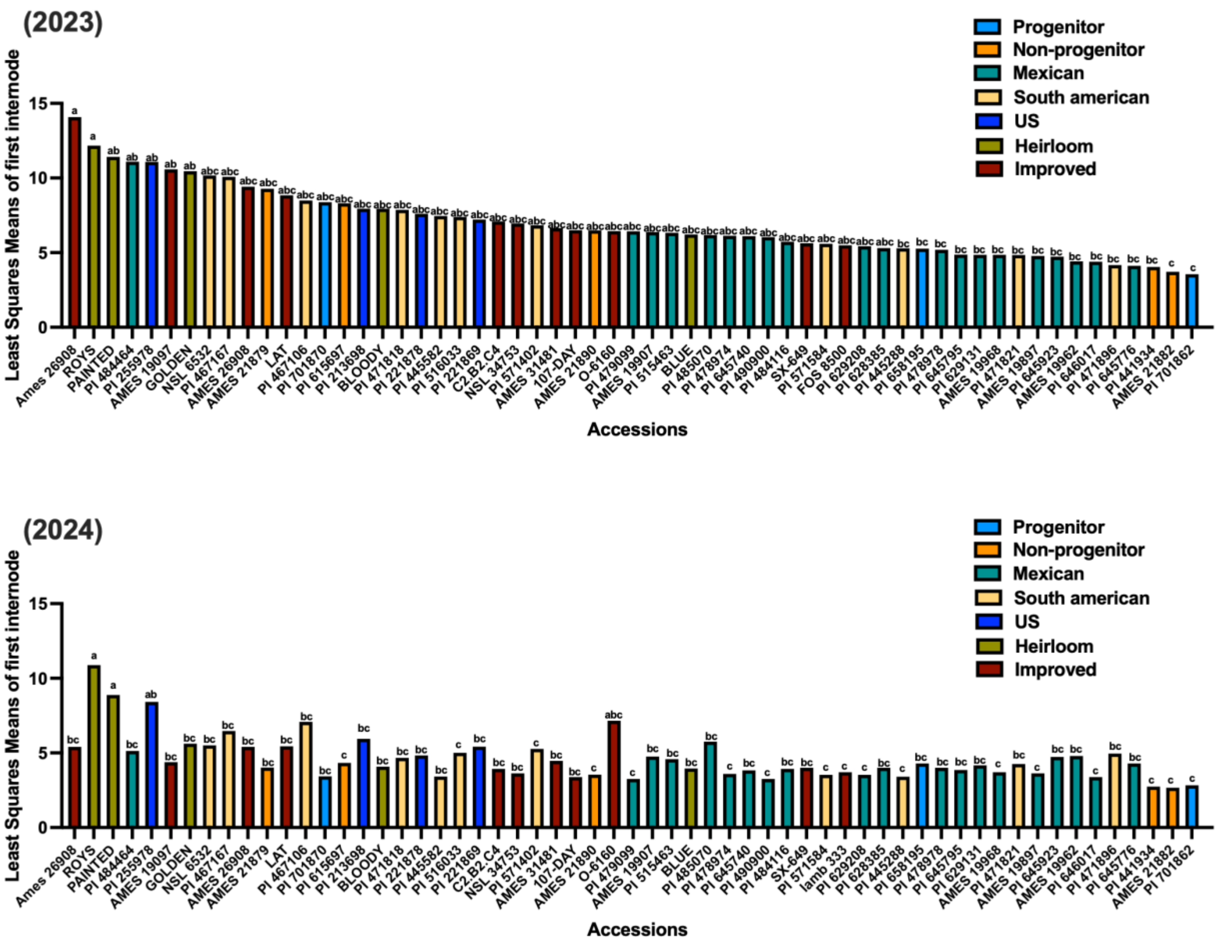


**Figure S5.** Least-squares means of first internode across maize accessions representing different domestication groups (Progenitor, Non-progenitor, Mexican landraces, South American landraces, U.S. landraces, Heirloom, and Improved lines). Bars are colored by domestication group and ordered by descending mean ARLN. The panels show accessions without grouping letters and Tukey–Kramer multiple comparison groupings (α = 0.05); accessions sharing the same letter are not significantly different. Least-squares means were estimated from a generalized linear mixed model with a negative binomial distribution and log link, accounting for block as a random effect.

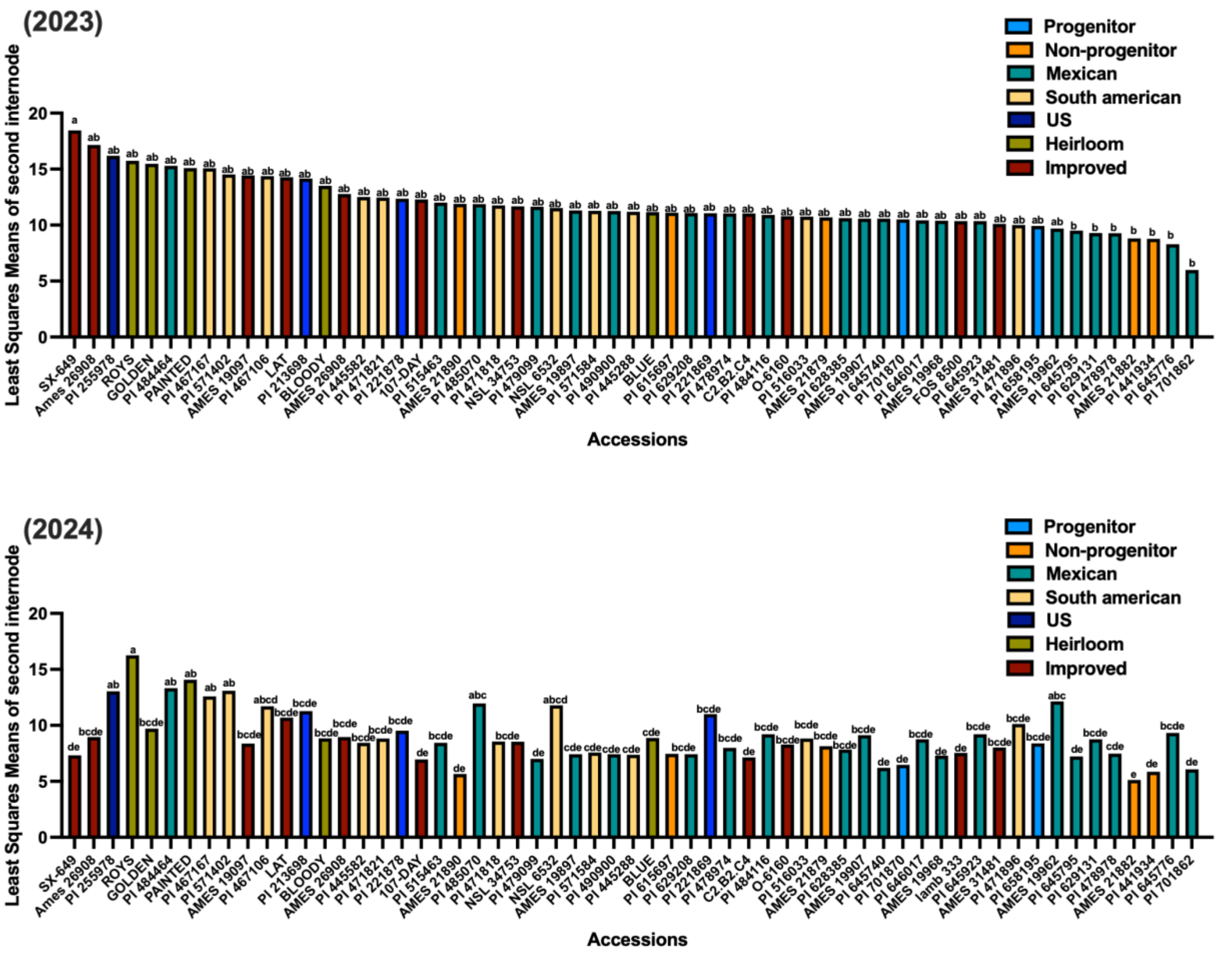


**Figure S6.** Least-squares means of second internode across maize accessions representing different domestication groups (Progenitor, Non-progenitor, Mexican landraces, South American landraces, U.S. landraces, Heirloom, and Improved lines). Bars are colored by domestication group and ordered by descending mean ARLN. The panels show accessions without grouping letters and Tukey–Kramer multiple comparison groupings (α = 0.05); accessions sharing the same letter are not significantly different. Least-squares means were estimated from a generalized linear mixed model with a negative binomial distribution and log link, accounting for block as a random effect.

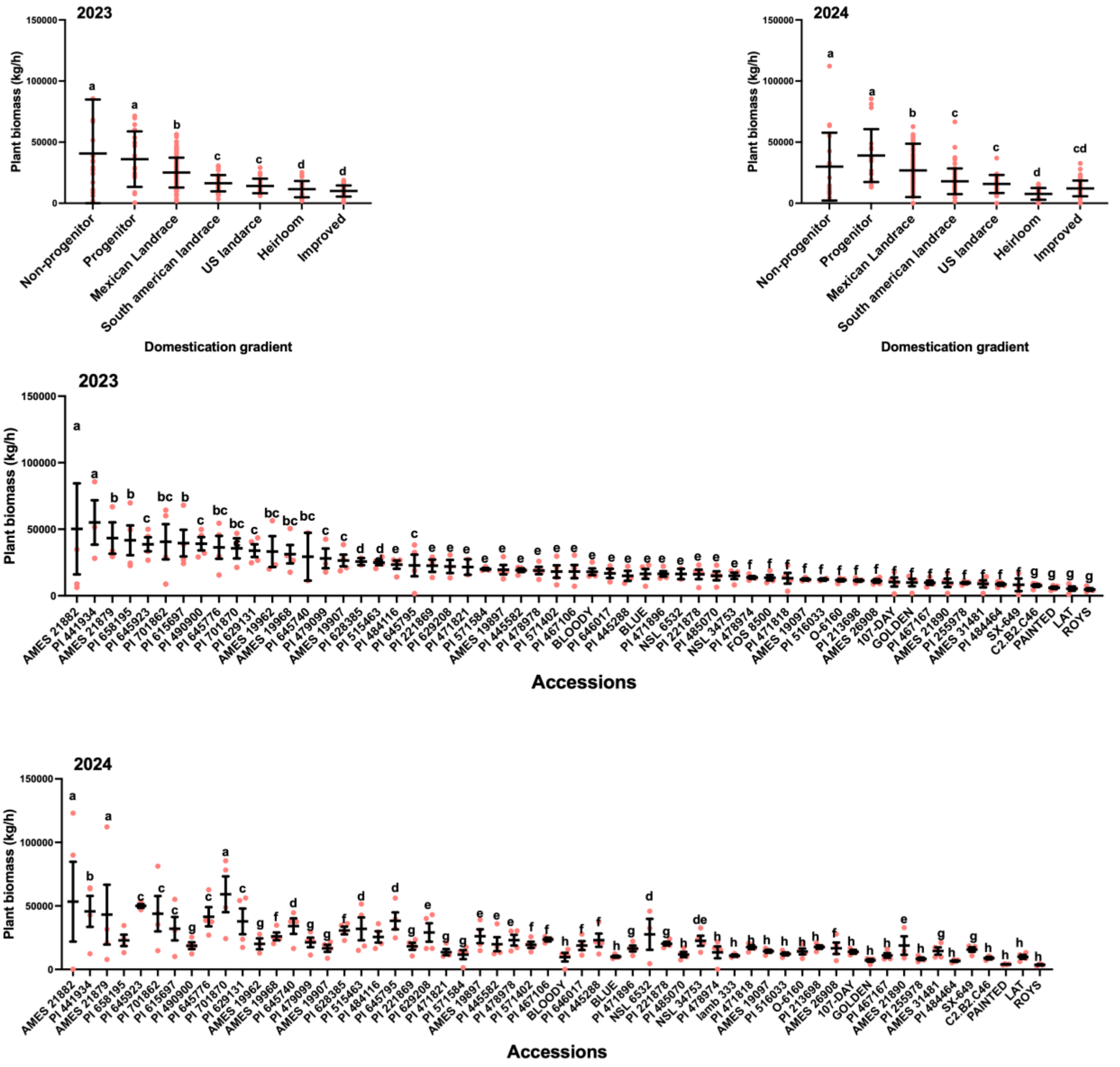


**Figure S7.** Least-squares means of plant biomass across maize accessions representing different domestication groups (Progenitor, Non-progenitor, Mexican landraces, South American landraces, U.S. landraces, Heirloom, and Improved lines). The panels show accessions without grouping letters and Tukey–Kramer multiple comparison groupings ($\alpha = 0.05$); accessions sharing the same letter are not significantly different. Least-squares means were estimated from a generalized linear mixed model with a negative binomial distribution and log link, accounting for block as a random effect.

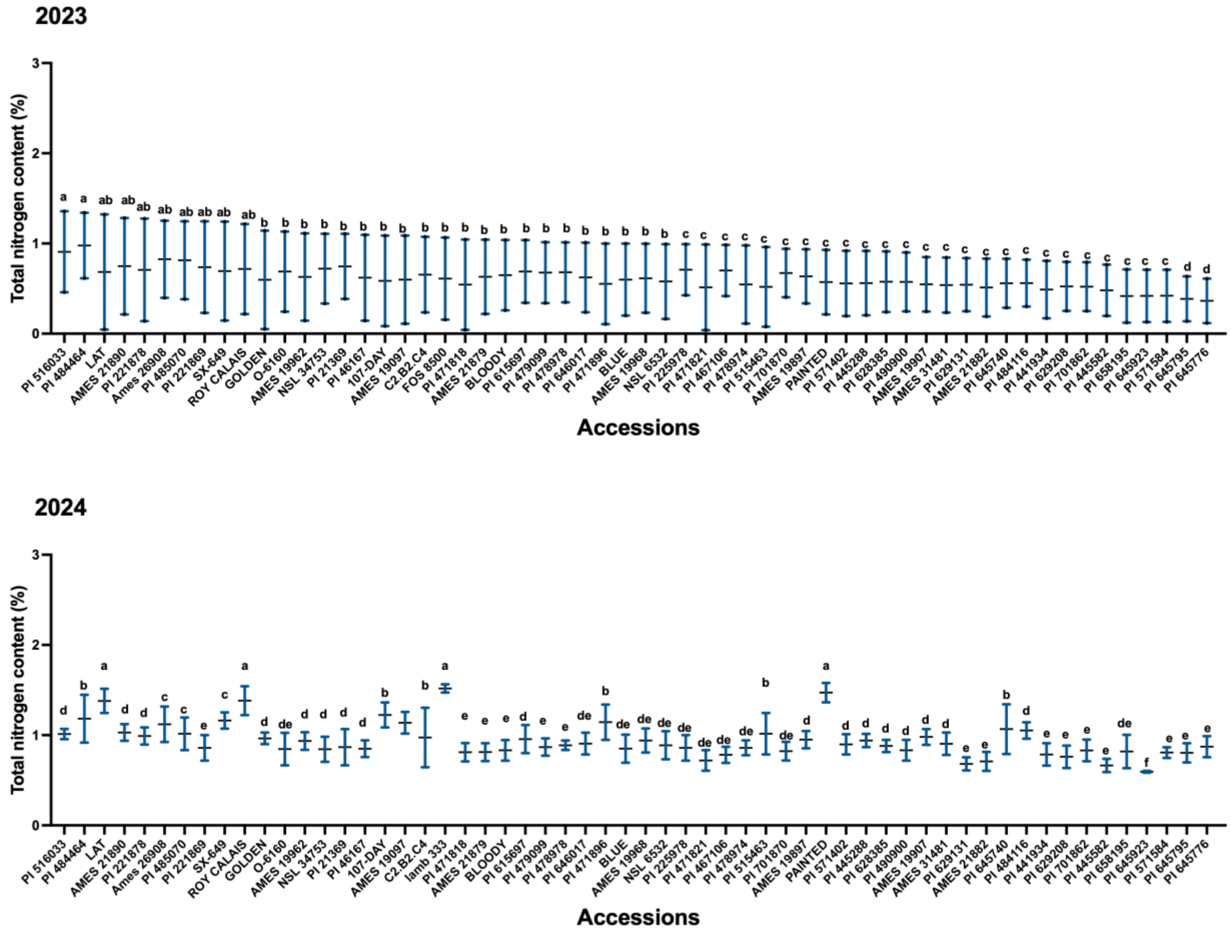


**Figure S8.** Variation in total nitrogen content among accessions in 2023 and 2024. Accessions represent stages along the domestication gradient: non-progenitor taxa, progenitors, Mexican landraces, South American landraces, U.S. landraces, heirloom varieties, and improved lines. Boxplots show the distribution of observations within each group, with central lines indicating medians and whiskers representing variability. Letters above groups indicate significant differences among domestication categories based on post hoc multiple comparisons ($P < 0.05$). The results were derived using generalized linear mixed models (GLMMs) fitted, accounting for block as random effects and accession as fixed effect.

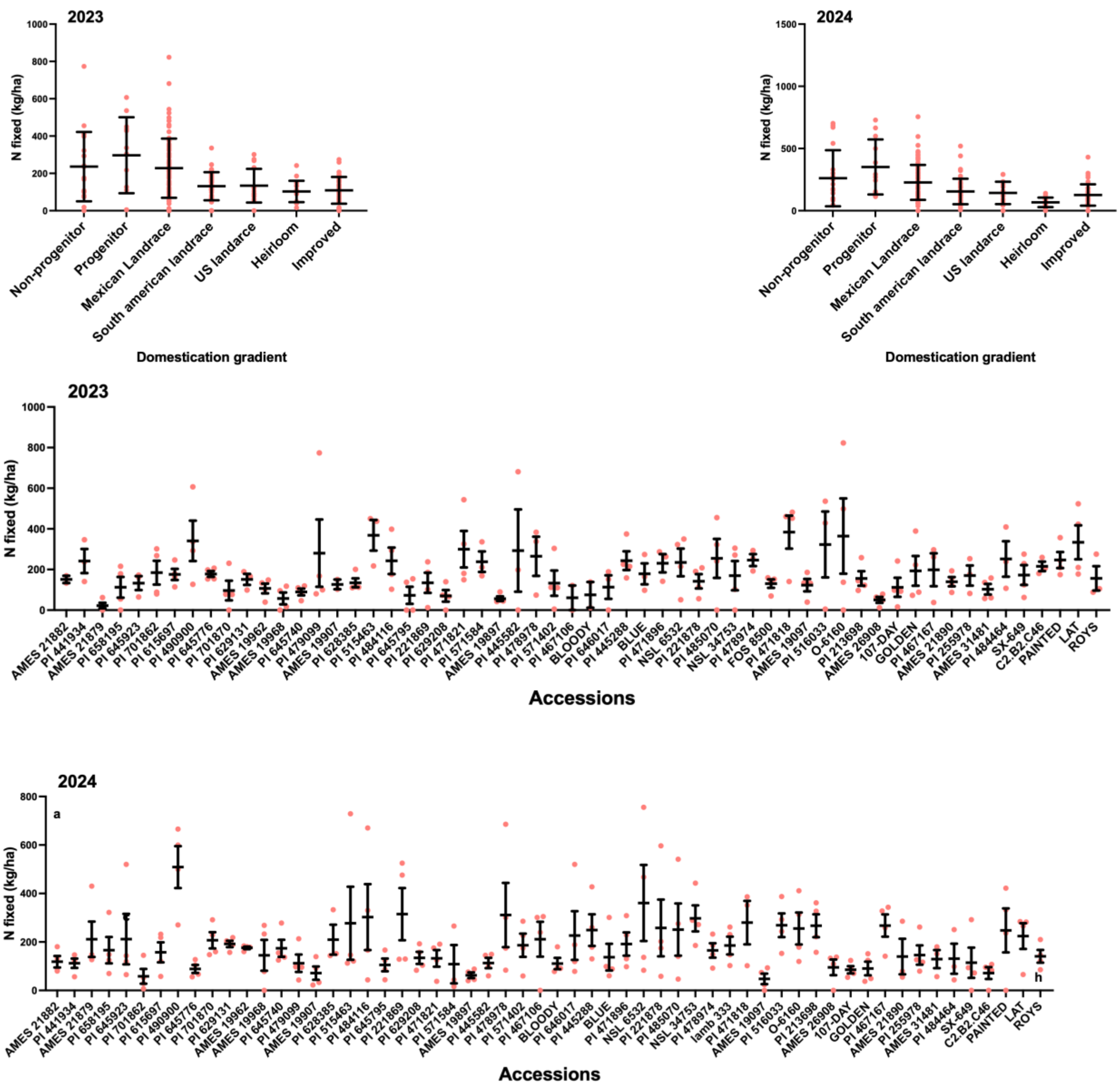


**Figure S9.** Least-squares means of nitrogen fixed (kg/ha) across maize accessions representing different domestication groups (Progenitor, Non-progenitor, Mexican landraces, South American landraces, U.S. landraces, Heirloom, and Improved lines). The panels show accessions without grouping letters and Tukey–Kramer multiple comparison groupings (α = 0.05); accessions sharing the same letter are not significantly different. Least-squares means were estimated from a generalized linear mixed model with a negative binomial distribution and log link, accounting for block as a random effect.

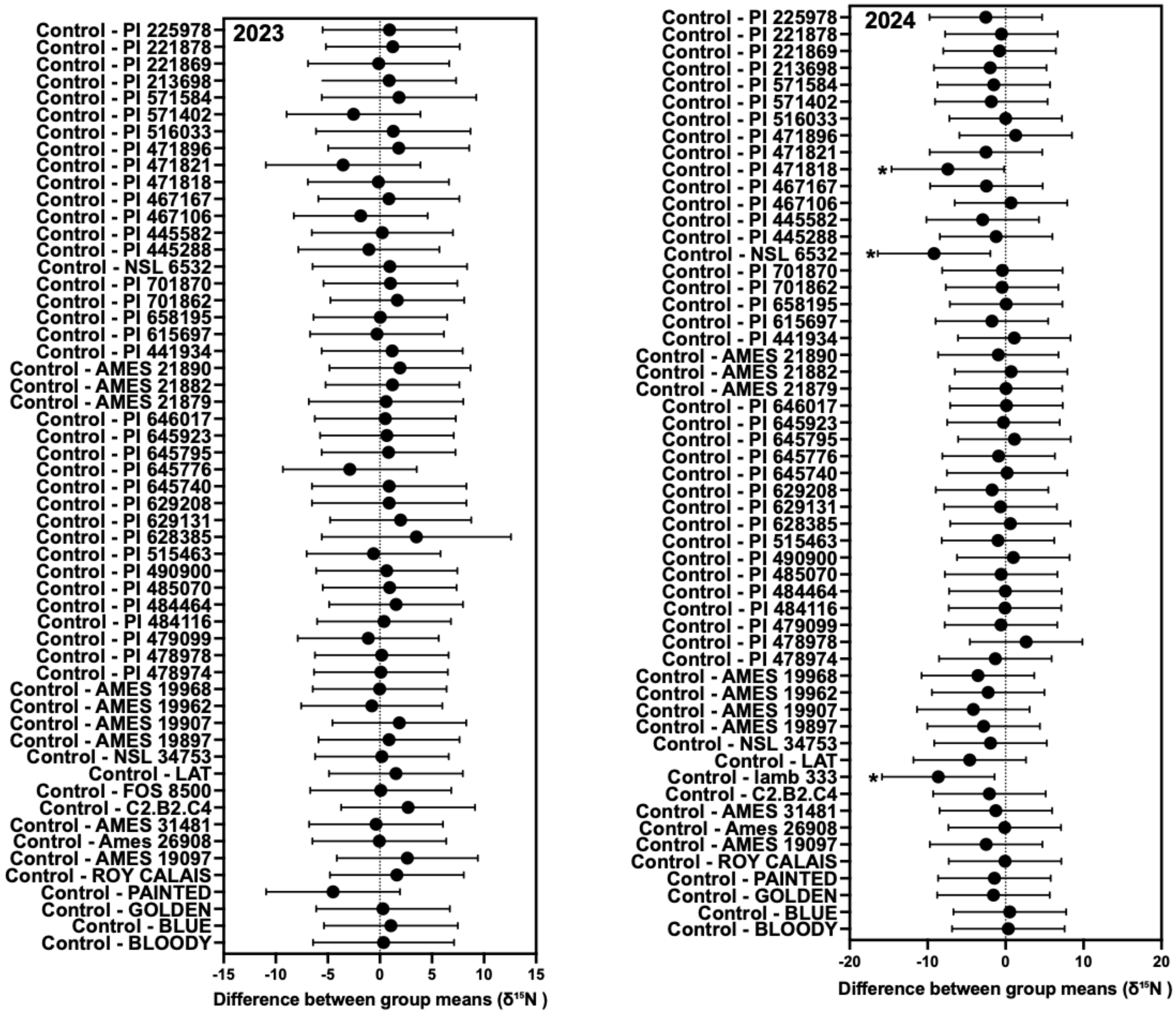


**Figure S10.** Figure S10. Comparison of $\delta^{15}N$ values between Midwest reference varieties and individual Zea accessions using Dunnett's multiple-comparison test. Estimated differences in least-squares means (LSMeans) of $\delta^{15}N$ are shown for the 2023 (left) and 2024 (right) field experiments. Contrasts are expressed as the Midwest reference group minus each accession (Control − accession), as indicated by the contrast labels. Thus, a negative difference indicates that the accession had a higher $\delta^{15}N$ value than the Midwest reference, whereas a positive difference indicates that the accession had a lower $\delta^{15}N$ value than the reference. Because lower plant $\delta^{15}N$ relative to a non-fixing reference is consistent with greater atmospheric N acquisition, positive—not negative—contrasts are in the direction expected with greater N fixation relative to the Midwest reference. Points represent estimated differences in LSMeans, horizontal bars indicate 95% confidence intervals, and the vertical dashed line denotes no difference between an accession and the reference group (difference = 0). Asterisks denote accessions that differed significantly from the reference group according to Dunnett-adjusted comparisons ($P < 0.05$).

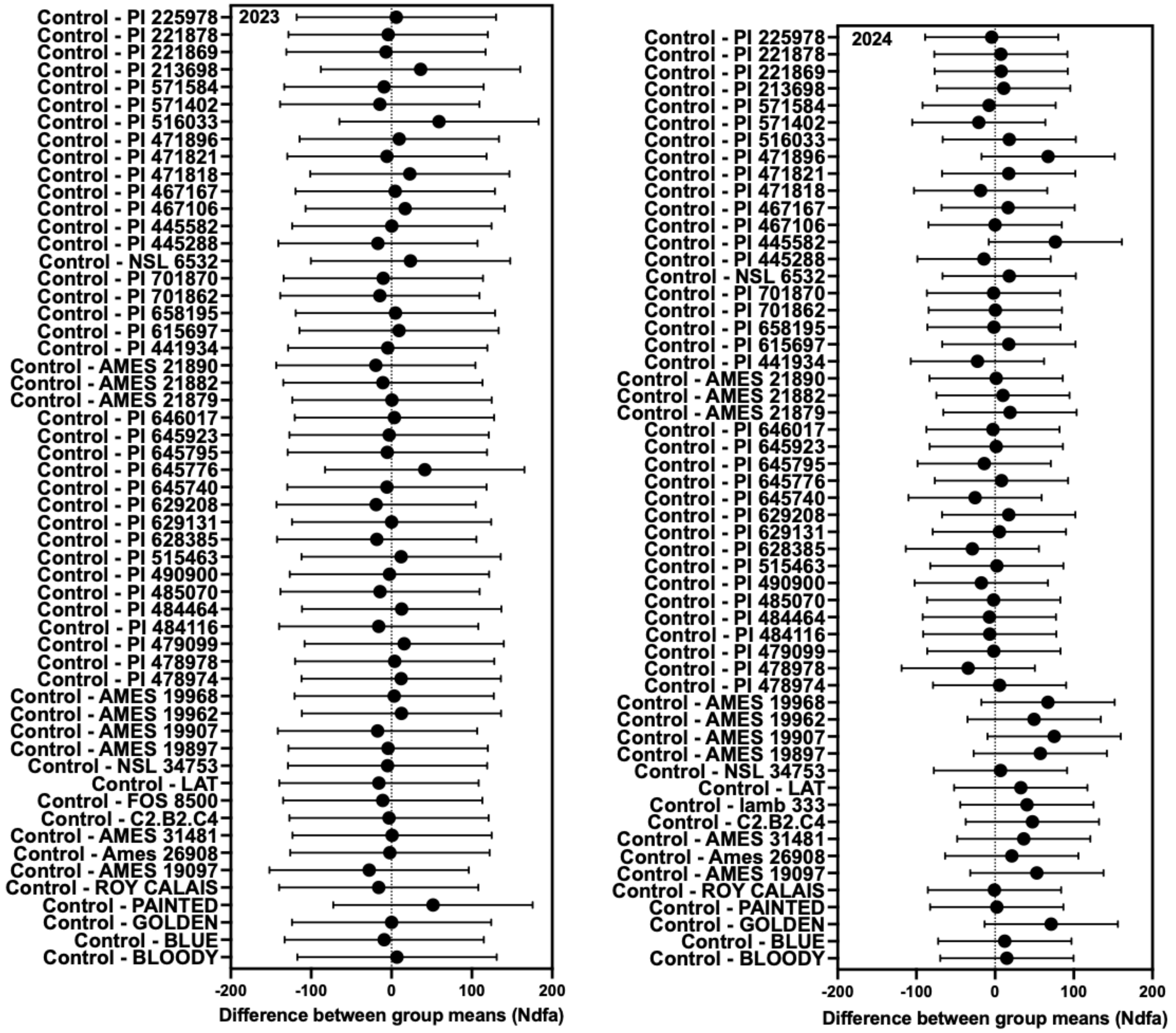


**Figure S11.** Comparison of Ndfa values between Midwest reference varieties and individual Zea accessions using Dunnett's multiple-comparison test. Estimated differences in least-squares means (LSMeans) of $\delta^{15}N$ between each accession and the Midwest reference group (Control) are shown for the 2023 (left) and 2024 (right) field experiments. Points represent estimated mean differences, and horizontal bars indicate 95% confidence intervals. The vertical dashed line denotes no difference between an accession and the reference group (difference = 0). Negative values indicate lower $\delta^{15}N$ than the Midwest reference, consistent with a greater proportion of nitrogen derived from atmospheric fixation, whereas positive values indicate higher $\delta^{15}N$. Asterisks denote accessions that differed significantly from the reference group according to Dunnett-adjusted comparisons ($P < 0.05$).

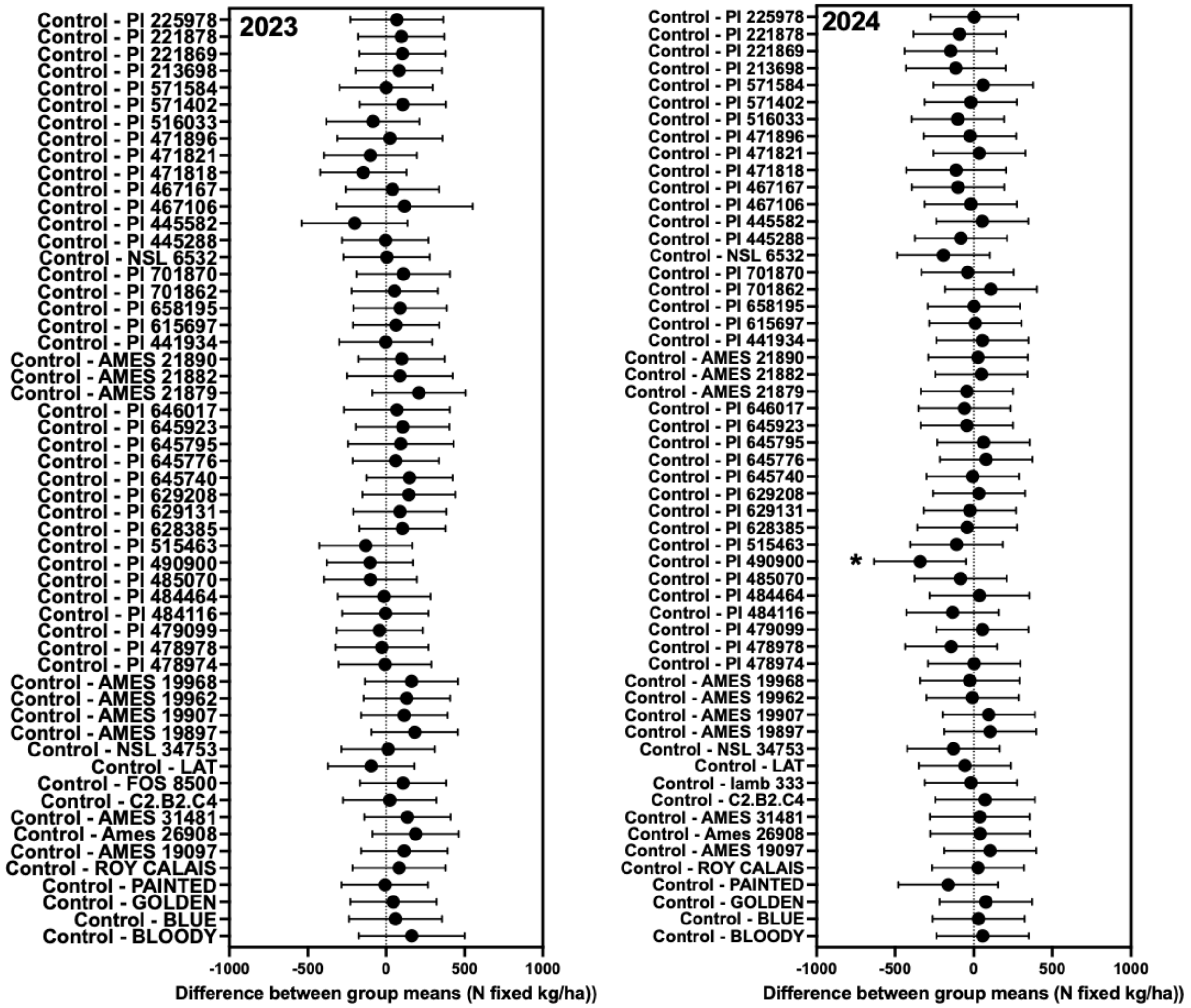


**Figure S12.** Comparison of N fixed kg/ha values between Midwest reference varieties and individual Zea accessions using Dunnett's multiple-comparison test. Estimated differences in least-squares means (LSMeans) of $\delta^{15}N$ between each accession and the Midwest reference group (Control) are shown for the 2023 (left) and 2024 (right) field experiments. Points represent estimated mean differences, and horizontal bars indicate 95% confidence intervals. The vertical dashed line denotes no difference between an accession and the reference group (difference = 0). Negative values indicate lower $\delta^{15}N$ than the Midwest reference, consistent with a greater proportion of nitrogen derived from atmospheric fixation, whereas positive values indicate higher $\delta^{15}N$. Asterisks denote accessions that differed significantly from the reference group according to Dunnett-adjusted comparisons ($P < 0.05$).

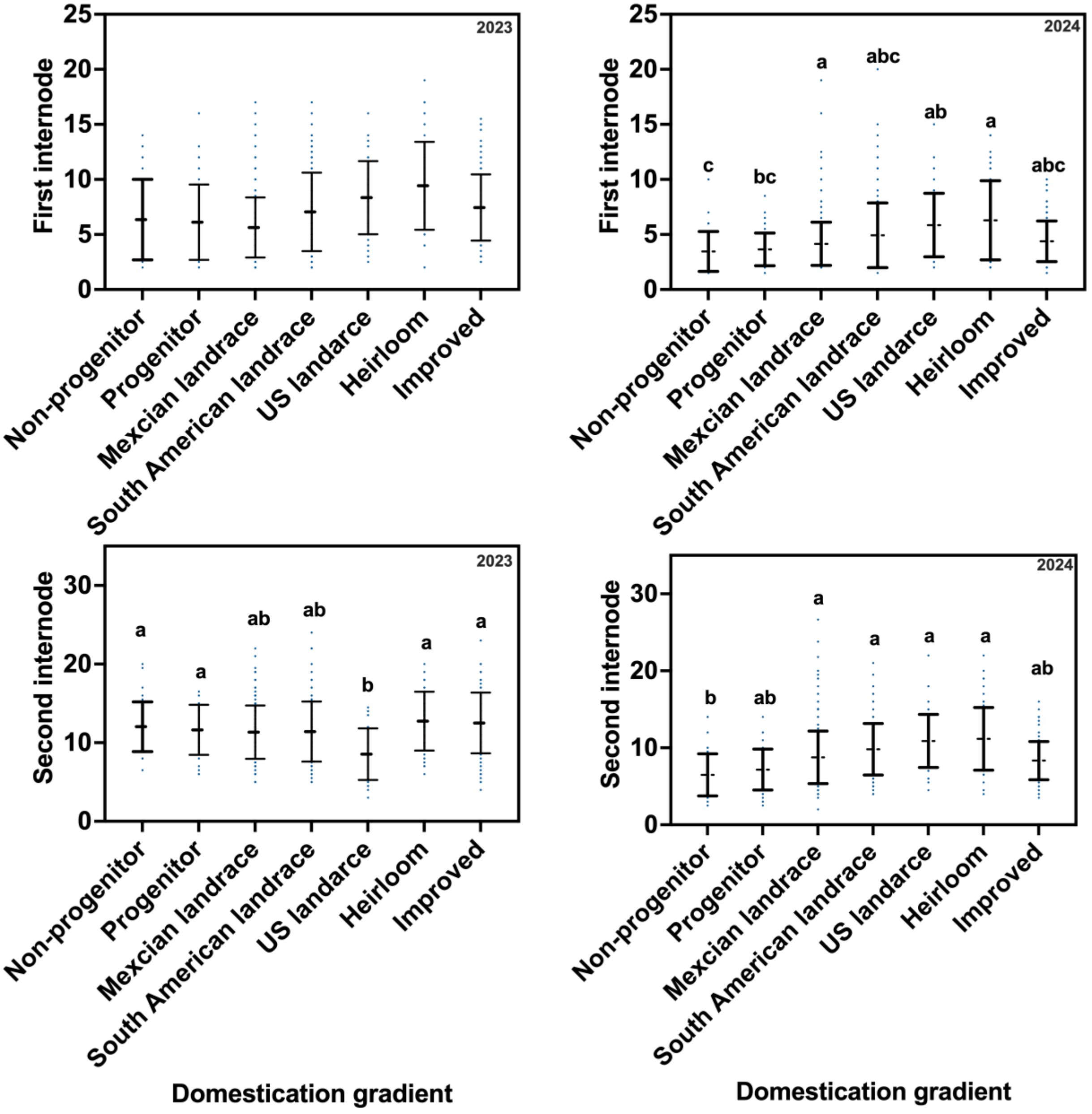


**Figure S13.** Variation in first and second internodes across the *Zea* domestication gradient in two consecutive years (2023 and 2024). Panels show LS means (± SE) and individual domestication values forAccessions are grouped into eight domestication categories: Non-progenitor Teosinte, Progenitor Teosinte, Mexican landrace, South American landrace, US landrace, Heirloom, and Improved. Different letters above bars indicate significant differences among domestication groups (Tukey's adjustment, $P < 0.001$).

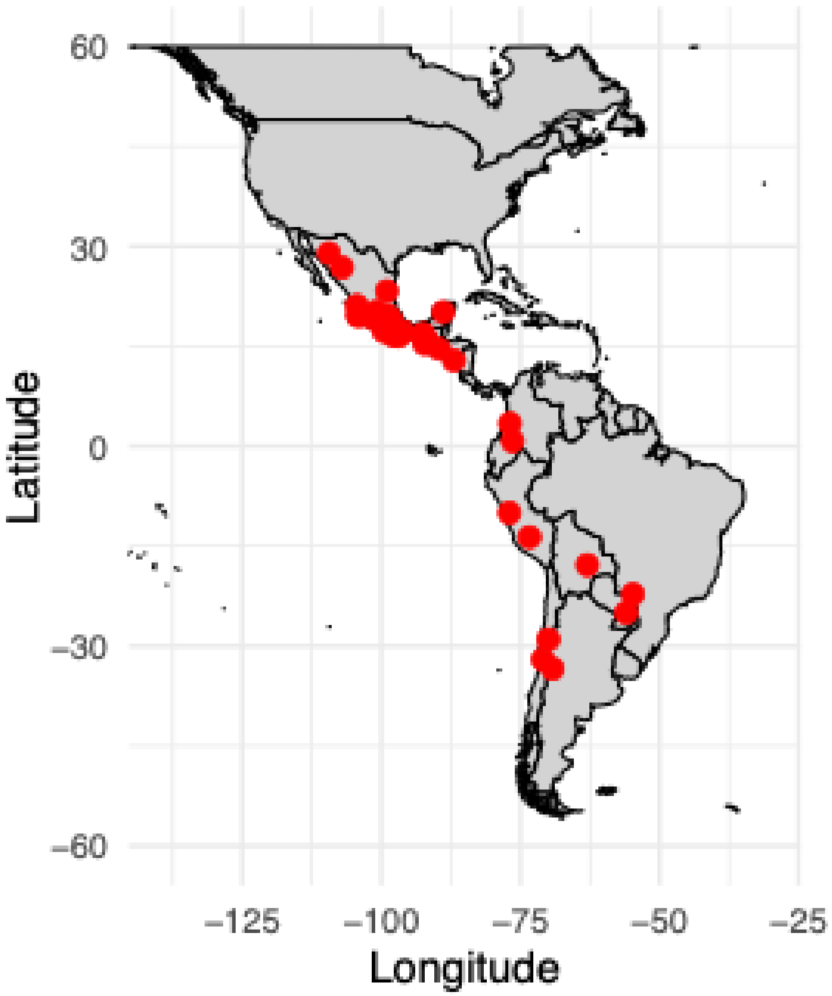


**Figure S14.** Geographic location of teosinte, Mexican and South American landraces within the center of origin.
